# Genetic regulation of fatty acid content in adipose tissue

**DOI:** 10.1101/2025.05.28.655990

**Authors:** Xinyu Yan, Amy L. Roberts, Julia S. El-Sayed Moustafa, Sergio Villicaña, Maryam Al-Hilal, Max Tomlinson, Cristina Menni, Thomas A. B. Sanders, Maxim B. Freidin, Jordana T. Bell, Kerrin S. Small

## Abstract

Fatty acids are important as structural components, energy sources, and signaling mediators. While studies have extensively explored genetic regulation of fatty acids in serum and other bodily fluids, their regulation within adipose tissue, a crucial regulator of cardiovascular and metabolic health remains unclear. Here, we investigated the genetic regulation of 18 fatty acids in subcutaneous adipose tissue from 569 female twins from TwinsUK. Using twin models, the heritability of fatty acids ranged from 5% to 59%, indicating a substantial genetic regulation of fatty acid levels within adipose tissue, which was also tissue-specific in many cases. Genome-wide association studies identified ten significant loci, in *SCD, ADAMTSL1, ZBTB41, SNTB1, EXOC6B, ACSL3, LINC02055, MKRN2/TSEN2, FADS1* and *HAPLN* across 13 fatty acids or fatty acid product-to-precursor ratios. Using adipose gene expression and methylation, which were concurrently measured in these samples, we detected five fatty acid-associated signals that colocalized with eQTL and meQTL signals, highlighting fatty acids that are regulated by molecular processes within adipose tissue. We identified strong associations of adipose fatty acids-associated loci with type 2 diabetes, body fat percentage, and cardiovascular disease. We explored links between polygenic scores of common metabolic traits and adipose fatty acid levels, and identified associations between polygenic scores of BMI, body-fat distribution and triglycerides and several fatty acids, indicating these risk scores impact local adipose tissue content. Overall, our results identified local genetic regulation of fatty acids within adipose tissue and highlighted their links with renal and cardio-metabolic health.

## Introduction

Adipose tissue is a major endocrine organ and a crucial regulator of cardiovascular health, secreting a range of bioactive products such as adipokines, leptin, adiponectin, and other hormones through endocrine and paracrine mechanisms, and it lies at the heart of a regulatory network influencing energy balance, glycolipid metabolism, cardio-metabolic health, and immune response ^1–3^. Adipose tissue is the largest fat depot, storing and releasing fatty acids as needed ^4^. Fatty acids are monocarboxylic acids with a long hydrocarbon linear chain, saturated or unsaturated acids, with an even number of carbon atoms. Fatty acids function as energy sources, structural components, and signaling mediators, and contribute to a range of molecular processes critical to common diseases, including neurological and cardiovascular disorders, type 2 diabetes (T2D), and cancers ^2,5^. Understanding the genetic regulation of fatty acids in adipose tissue may reveal the genetic and molecular mechanisms underlying risk of cardio-metabolic and other disorders.

Due to the fundamental role of circulating fatty acids, and accessibility of blood samples, a plethora of studies have investigated the genetic regulation of circulating fatty acids, usually profiled in serum or plasma. Circulating fatty acids are under strong genetic regulation, with heritability estimates ranging from 10% to 48% ^6,7^. To date, genome-wide association studies (GWASs) have identified thousands of single-nucleotide polymorphisms (SNPs) associated with fatty acid metabolism in plasma or serum (GWAS Catalog, https://www.ebi.ac.uk/gwas/), and several recent studies have conducted GWAS of fatty acids in bodily fluids such as faeces ^8^, urine ^9^ and saliva ^10,11^. In contrast, limited studies ^12,13^ have explored genetic associations of adipose tissue fatty acids but these are limited to candidate SNP studies or GWAS limited to estimates of three fatty acid desaturases activity [delta-6-desaturase (D5D), delta-6-desaturase (D6D) and stearoyl-CoA desaturase-1 (SCD)] ^13^, and there are currently no publicly accessible GWAS summary statistics of adipose fatty acid levels. Here, we investigate the genetic regulation of adipose tissue fatty acids levels by profiling 18 adipose fatty acids in subcutaneous adipose tissue biopsies from 569 females from the TwinsUK cohort. We utilize classic twin models to estimate heritability of these fatty acids and their ratios (fatty acid ratios can serve as proxies of enzymatic activity), and compare them with matched serum data from the same individuals to estimate tissue specificity of heritability. We conducted GWASs for the 18 fatty acids and 15 heritable fatty acid ratios and identified ten genome-wide significant association signals. To elucidate the underlying molecular mechanisms at the GWAS loci, we integrated the fatty acid levels with both eQTL and meQTL data and concurrently measured adipose gene expression and DNA methylation from the same samples. Finally, to define the link between genetic regulation of cardio-metabolic traits and adipose fatty acids, we performed phenome-wide association studies of GWAS lead variants and explored the association between polygenic scores of cardio-metabolic traits and fatty acid levels. Overall, our results provide insights into the regulation of these key molecules within a metabolic tissue.

## Material and Methods

### Participants and samples

The study included 569 female twins from the TwinsUK population cohort ^14^ with available adipose fatty acid data. The study cohort consisted of 243 twin pairs (105 monozygotic (MZ) and 138 dizygotic (DZ) twin pairs) and 83 singletons of European ancestry with a mean age of 59 (SD 9) years old and a mean body mass index (BMI) of 26.53 (SD 4.67) kg/m^2^. Punch biopsies were taken from a sunlight-protected area of the stomach adjacent and inferior to the umbilicus, and immediately divided into aliquots and snap frozen, with one aliquot allocated to fatty acid profiling, one for RNA extraction, and one for DNA extraction and methylation profiling. Serum samples were collected at the same time point as the adipose tissue during clinical visits. All samples and information were collected with written and signed informed consent, including consent to publish within the TwinsUK study. TwinsUK has received ethical approval associated with TwinsUK Biobank (19/NW/0187), TwinsUK (EC04/015) or Healthy Ageing Twin Study (HATS) (07/H0802/84) studies from NHS Research Ethics Service Committees London – Westminster. The study design is presented in Figure 1.

**Figure 1:**
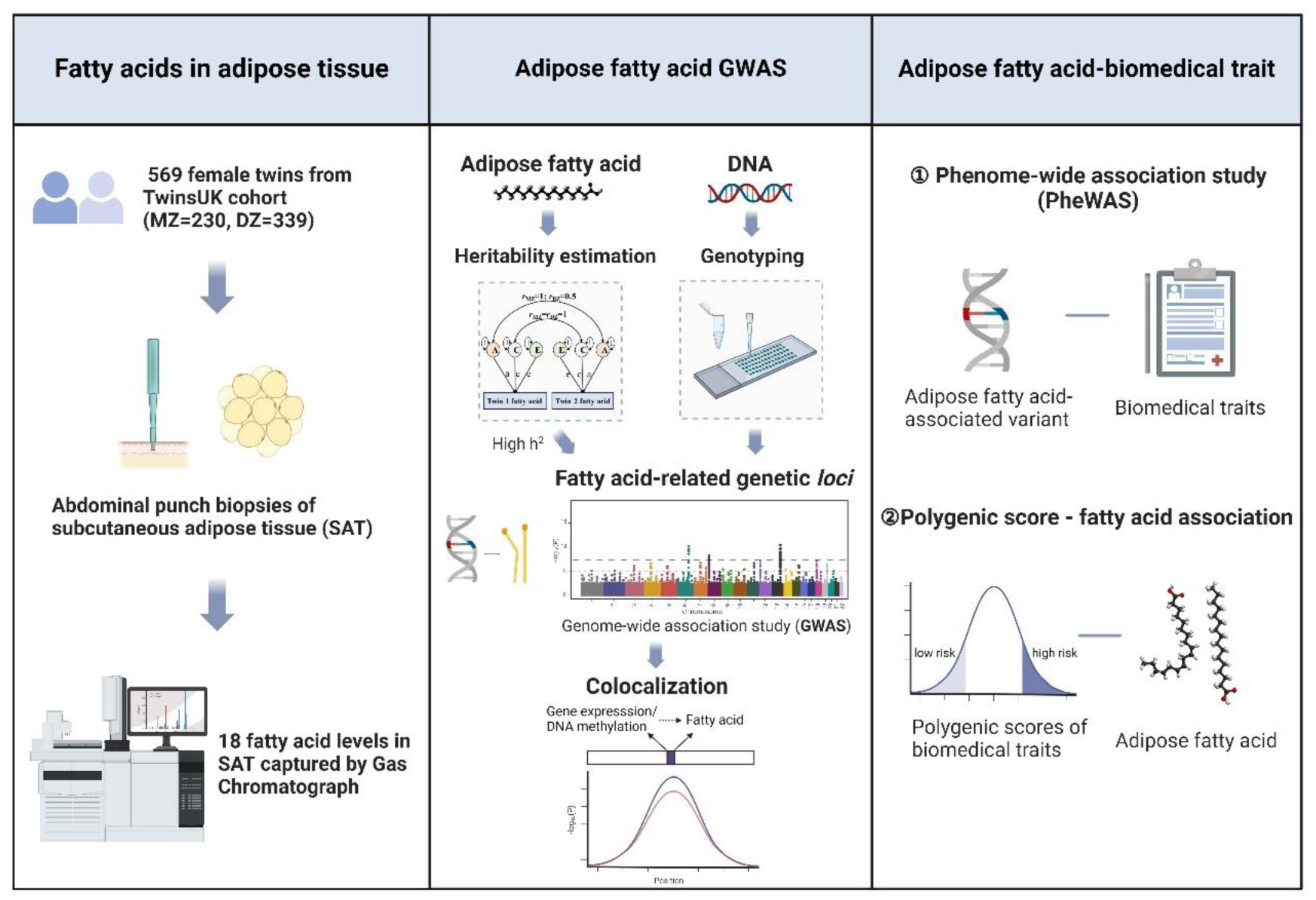
Study design of genetic regulation of fatty acid content in adipose tissue.

### Fatty acid profiling

#### Adipose fatty acid profiling

Fatty acid methyl esters were separated on an Agilent 7890 Gas Chromatograph (Agilent Technologies) fitted with a 60 m capillary column BPX70 with hydrogen as carrier gas and a flame ionization detector. The identities of fatty acids for which standards were not available were estimated using plots of the retention times as described elsewhere ^15^ to obtain the equivalent chain lengths and n-9/n-6/n-3 ratios. Quality control was maintained by running a matrix standard of the sample of adipose tissue fatty acid methyl esters with each run. Chromatograms were evaluated using ChemStation software (version 8.04) and the output was exported via a visual basic application into Excel for data analysis. Detailed information was described elsewhere ^16^. A total of 18 types of fatty acids were quantified as proportions to the total fatty acid. Adipose fatty acids were summed up to obtain three classes of fatty acid values for each participant, saturated fatty acid (SFA), monounsaturated fatty acid (MUFA), and polyunsaturated fatty acid (PUFA). Adipose fatty acid ratios were calculated as product-to-precursor ratios of fatty acids to reflect the conversion rate of fatty acids ^17^. Rank-based inverse normal transformation was applied to the raw adipose fatty acid and fatty acid ratio values before analyses.

#### Serum metabolite profiling

Fatty acids in serum were extracted from the metabolite dataset in TwinsUK which includes 15 fatty acids. The non-targeted metabolomics analysis was performed at Metabolon (Durham, North Carolina, USA) on an ultra-high-performance liquid chromatography-tandem mass spectrometry (UPLC-MS/MS) instrument. Methods to process samples and data were described by Long et al ^7^. Date matched metabolon profiles were available for 471 twins. Free fatty acid levels were rank-based inverse normalized before analyzing.

### Genotyping and imputation

SNPs were genotyped using a combination of Illumina HumanHap300 (317k), HumanHap610Q (610k), 1M-Duo, and 1.2M-Duo (1M) arrays. Autosomal genotype data were imputed using the Haplotype Reference Consortium (HRC) v1.1 panel on the Michigan Imputation server (human reference build hg37). Quality control measures were applied, including Hardy-Weinberg equilibrium (HWE) *P* > 10^−6^, minimum genotyping success rate > 0.95, imputation quality R^2^ > 0.5, and minimum minor allele frequency (MAF) > 0.01, resulting in 7,656,749 retained autosomal SNPs. Human reference hg38 coordinates were generated using LiftOver. A total of 538 individuals had both autosomal genotype and fatty acid data.

Chromosome X genotypes were derived from whole genome sequencing as part of the UK10K project. Briefly, low-coverage whole genome sequencing data were generated using Illumina HiSeq 2000 and reads were mapped to human reference build hg37. Quality control measures were applied, including a 99.50 truth sensitivity threshold for variant filtering, HWE *P* > 10^−6^, and MAF > 0.01, resulting in 331,002 retained chromosome X SNPs. Human reference hg38 coordinates were generated using LiftOver. A total of 352 individuals had both chromosome X genotype and fatty acid data.

### Heritability estimates

Heritability of fatty acids in adipose tissue and serum in the TwinsUK cohort were estimated as proportions of total variance explained by additive genetic effects using the ACE twin model which modeled trait variance as a function of additive genetics (A), common environment (C) and unique environment effects (E) using the structural equation modeling software ‘OpenMx’ ^18^. In the model to estimate the heritability of fatty acids in adipose tissue, age was included as a covariate, followed by sensitivity analyses where (1) age and cell types [adipocyte, macrophage, and microvascular endothelial cell (MVEC)] ^19^; (2) age and BMI; (3) age, BMI and cell types were included as covariates. Only age was adjusted for when estimating the heritability of serum fatty acids.

### Discovery genome-wide association analysis

Discovery GWASs were performed across 18 individual fatty acids, three fatty acid sums (SFA, MUFA, and PUFA), and 15 heritable (h^2^ ≥ 0.15) fatty acid ratios of product-to-precursor fatty acids. The Wald test implemented in GEMMA V0.98.1 ^20^ was utilized to assess the significance of the associations, accounting for family relatedness using a sample kinship matrix, using linear mixed models adjusting for age. Three thresholds were applied: 1) Genome-wide significance: *P_G_* < 5 × 10^−8^; 2) Bonferroni-corrected study-wide genome-wide significance: *Pb* < 2.8 × 10^−9^ (5 × 10^−8^ / 18, controlled for the number of independent GWASs, the effective number of fatty acid (ratio) was derived from the ‘meff’ function in the R package ‘poolr’ by extracting the eigenvalues from a correlation matrix; 3) Suggestive significance: *Ps* < 10^−6^. To test whether any of the associated loci had multiple distinct association signals, conditional analysis was performed by including the lead variant (*P* < 5 × 10^−8^) as a covariate. The locus-wide significance threshold was an arbitrary set of *P* < 10^−4^ to identify secondary association signals within ±100 kb of the primary

GWAS signal. This threshold accounts for the reduced multiple testing burden due to local linkage disequilibrium (LD) structure, and balances sensitivity for detecting true secondary signals while controlling multiple testing within a defined locus. We also performed sensitivity analyses adjusted for (1) age and cell type composition; (2) age and BMI; (3) age, BMI and cell type composition in the GWAS model. The variance explained by the lead variant was calculated using the formula 2*EAF*(1-EAF)*Beta^2^/Var(Y), in which EAF is the effect allele frequency, Beta is the effect estimate of the effect allele, Var(Y) is the variance of the phenotype Y ^21^. Genomic inflation factor (lambda genomic control) was calculated using Devlin and Roeder’ formula ^22^. Manhattan plots and QQ plots were produced by the R package ‘qqman’ and Locus zoom plots were generated using the open-source software, LocusZoom.js.^23^.

### Replication genome-wide association analysis

#### Blood metabolites studies

(1) TwinsUK matched serum dataset and TwinsUK-KORA: Serum metabolites profiling method has been mentioned as above. We tested seven out of 17 SNP-fatty acid association pairs across five available individual fatty acids and five loci in the TwinsUK concurrent serum metabolite dataset (N = 448). Meta-analysis combining TwinsUK and KORA (Augsburg) participants’ metabolites GWAS results were further included (N = 7,824) ^24^. (2) EPIC-Norfolk and INTERVAL: Blood metabolites were measured using the untargeted platform Metabolon in 19,994 individuals from UK-based studies, EPIC-Norfolk and INTERVAL, and GWASs of metabolites were performed ^25^. (3) CLSA: Blood metabolites were quantified using UPLC-MS/MS platform in 8,299 individuals from a Canada-based study, CLSA, and GWASs were performed across metabolites ^26^.

### RNA-seq and DNA methylation

#### RNA-seq

RNA sequencing was performed to measure gene expression levels in TwinsUK participants in adipose tissue (N = 765), skin (N = 706), LCLs (N = 804), and whole blood (N = 389), as previously reported ^27^. RNA-Seq reads were aligned to the GRCh37 (hg19) reference genome using STAR v2.4.0.1 ^28^ and then quality control was conducted ^29,30^. Gene-level counts were quantified using the ‘quan’ function from QTLtools ^31^ and Gencode v19 ^32^. RNA-Seq gene expression data were filtered to retain genes with five or more counts per million (CPMs) in ≥ 25% of individuals. Read counts were adjusted for each gene for the trimmed mean of M-values (TMM) ^33^ and were rank-based inverse normalized.

#### DNA methylation

The DNA was bisulfite-converted using the EZ DNA Methylation Kit (Zymo Research). The DNA methylation profiles of adipose tissue were conducted in 588 twins using Illumina Infinium HumanMethylation450 BeadChip (450K array) ^34^ to obtain 𝛽-values, which represent the proportion of methylated probes at a specific CpG site ^35^. The raw methylation data was processed, including background correction, dye-bias adjustment, signal normalization, and the estimation of adjusted 𝛽-values, as described previously ^36^. There were 442,160 CpG sites of 539 participants who passed quality control checks and were included in downstream analyses. DNA methylation levels of CpG sites were rank-based inverse normalized prior to association analysis using linear models.

#### QTL Analysis

(1) Expression quantitative trait loci (eQTL): Association between SNP and gene expression level was performed using a linear mixed-effect regression model, adjusting for BMI, family, zygosity, SNP genotyping chip and PEER factors, following the methods described previously ^37,38^. Conditional eQTLs were carried out if the *P* value of the top SNP was lower than 10^−5^ by including the genotype of the top SNP as an additional covariate. (2) eQTL and sQTL across tissues: We used multi-tissue eQTL and sQTL summary statistics in ∼706 individuals which were generated from the GTEx project (v8, European ancestry) ^39^. (3) Methylation quantitative trait loci (meQTL): Association between SNP and DNA methylation level was performed adjusting for technical covariates, age, predicted smoking, family relatedness, genetic PCs, and non-genetic DNA methylation PCs from DNA methylation, as described previously ^40^.

### QTL-fatty acid colocalization analysis

#### eQTL/meQTL-fatty acid colocalization

To test whether the GWAS lead variant was colocalized with the lead variant of gene expression or DNA methylation level, we filtered pairwise LD r^2^ between the GWAS lead variant and eQTL/meQTL lead variant within 1Mb greater than 0.6 of European ancestry population using the ‘LDpair’ module on ‘NIH LDlink’ (https://ldlink.nci.nih.gov/?tab=apiaccess). Next, to formally test if gene expression/DNA methylation (trait 1) and adipose fatty acid (trait 2) share a common genetic causal variant in a given region (hypothesis 4, shorted for H4), we ran GWAS-eQTL/meQTL colocalization analysis within 500Kb using ‘coloc’ v5.1.0.1 R package ^41^, followed by sensitivity analysis. We defined the variants as colocalized when the posterior probability of colocalized signal (PP.H4) was greater than 0.75 ^41^. We also colocalized GWAS secondary variant with eQTL/meQTL lead variant and colocalized GWAS lead variant with eQTL/meQTL secondary variant using the same pipeline.

#### Cell type composition estimates

Adipose tissue is comprised of multiple cell types, with the most common being adipocyte, endothelial and immune cells. We estimated adipose tissue cell type proportion from RNA-Seq data in TwinsUK, including adipocytes, macrovascular endothelial cells and macrophages, as previously described ^19^.

#### Fatty acid gene expression/DNA methylation association analysis

We assessed the association between adipose gene expression and fatty acids adjusting for technical covariates, age, BMI and family relatedness as covariates by linear mixed model using ‘lme4’ R package ^42^. Then we tested the association between adipose DNA methylation and fatty acids adjusting for technical covariates, age, predicted smoking, family relatedness, and non-genetic DNA methylation PCs. Gene expression or DNA methylation was treated as a continuous independent fixed effect, and each adipose fatty acid was treated as a continuous dependent variable.

### Fatty acid-kidney trait colocalization analysis

To test whether n-6 PUFAs in adipose tissue shared a common genetic causal variant with kidney traits, we integrated GWASs of TwinsUK adipose fatty acids with published GWASs of kidney trait ^43^ using the same pipeline. Both arachidonic acid level and the ratio of arachidonic acid to linoleic acid were included. Three kidney traits were included in the analysis: eGFR, serum creatinine and cystatin C.

### Phenome-wide association analysis

To identify whether adipose fatty acids-associated loci regulate other biomedical traits, we accessed phenome-wide association studies (PheWASs) of the lead variants from GWASATLAS (https://atlas.ctglab.nl/) and NIH Accelerating Medicines Partnership (AMP) (https://hugeamp.org/). The significant association was defined as *P* < 10^−5^ to account for multiple testing across ∼3302 phenotypes.

### Polygenic score analysis

We constructed polygenic scores (PGSs) for 13 common metabolic traits: abdominal subcutaneous adipose tissue volumes adjusted for BMI (ASATadjBMI), visceral adipose tissue volumes adjusted for BMI (VATadjBMI), gluteofemoral adipose tissue volumes adjusted for BMI (GFATadjBMI), BMI, waist-to-hip ratio adjusted for BMI (WHRadjBMI), type 1 diabetes (T1D), T2D, coronary artery disease (CAD), hypertension, high-density lipoproteins cholesterol (HDL), low-density lipoproteins cholesterol (LDL), log-transformed total triglycerides (TG) and total cholesterol (TC) ^44–50^. PGSs were calculated for 6847 participants using the linear scoring method implemented in PLINK 1.9 ‘--score’ function (Table S1). Then, we validated PGSs in TwinsUK where PGSs showed broadly strong associations with corresponding traits (Figure S1, Table S2 and Supplemental Materials and Methods). Finally, we tested the association between PGSs and fatty acid levels and summary levels of fatty acids (SFA, MUFA and PUFA), adjusting for age, cell types and relatedness excluding participants with T1D, T2D, cardiovascular disease (CVD) or hypertension. To overcome the multiple test burden, *P* values were Bonferroni-corrected using the effective number of fatty acids (N_eff_ = 13) for each PGS.

## Results

### Fatty acid characteristics in adipose tissue

We measured 18 types of fatty acids among 569 female twins (243 twin pairs and 83 singletons) in subcutaneous adipose tissue (Figure 1), including six saturated fatty acids (SFAs), four mono-unsaturated fatty acids (MUFAs), eight polyunsaturated fatty acids (PUFA) including four n-6 family and four n-3 family PUFA. Summary level values of SFA, MUFA and PUFA were calculated from individual measurements. The characteristics of adipose fatty acid proportions in our study are shown in Table 1. Overall, the values are in line with expectations from previous studies of adipose tissue ^51,52^, including high median levels of palmitic acid, oleic acid and linoleic acid, which together account for ∼78 percent of fatty acids within adipose tissue.

**Table 1.** Characteristics and heritability of fatty acid in adipose tissue.

| Adipose fatty acid | Common name | N | Median | Mean | Variance | Adipose fatty acid $h^2$ | Serum fatty acid $h^2$ |
| --- | --- | --- | --- | --- | --- | --- | --- |
| **SFA** | **Saturated fatty acid** | 543 | 25.62 | 25.74 | 8.28 | 0.28 | -- |
| C10:0 | Capric acid | 546 | 0.25 | 0.33 | 0.13 | 0 | 0.27 |
| C12:0 | Lauric acid | 566 | 0.34 | 0.61 | 0.53 | 0.09 | 0.05 |
| C14:0 | Myristic acid | 567 | 2.17 | 2.03 | 0.75 | 0 | 0.44 |
| C16:0 | Palmitic acid | 569 | 19.28 | 19.24 | 4.23 | 0.30 | 0.34 |
| C18:0 | Stearic acid | 569 | 3.10 | 3.21 | 0.89 | 0.51 | 0.33 |
| C20:0 | Arachidic acid | 563 | 0.35 | 0.36 | 0.03 | 0 | -- |
| **MUFA** | **Monounsaturated fatty acid** | 502 | 53.25 | 52.97 | 10.68 | 0.12 | -- |
| C16:1n-7 | Palmitoleic acid | 567 | 5.75 | 5.79 | 2.77 | 0.05 | 0.42 |
| C18:1n-7 | Vaccenic acid | 504 | 0.29 | 0.88 | 0.88 | 0.04 | -- |
| C18:1n-9 | Oleic acid | 569 | 46.77 | 46.55 | 9.29 | 0 | 0.40 |
| C18:1trans | Elaidic acid, trans-vaccenic acid | 497 | 0.51 | 0.53 | 0.09 | 0.43 | -- |
| **PUFA** | **Polyunsaturated fatty acid** | 557 | 14.79 | 15.03 | 6.06 | 0.31 | -- |
| C18:3n-3 | α-linolenic acid (ALA) | 564 | 0.85 | 0.85 | 0.05 | 0.50 | 0.21 |
| C20:5n-3 | Eicosapentaenoic acid (EPA) | 561 | 0.17 | 0.22 | 0.02 | 0 | 0.49 |
| C22:5n-3 | Docosapentaenoic acid (DPA) | 561 | 0.36 | 0.39 | 0.03 | 0.29 | 0.22 |
| C22:6n-3 | Docosahexaenoic acid (DHA) | 561 | 0.31 | 0.36 | 0.05 | 0 | 0.31 |
| C18:2n-6 | Linoleic acid (LA) | 568 | 11.92 | 12.05 | 5.61 | 0.42 | 0.31 |
| C20:3n-6 | Dihomo-γ-linolenic acid (DGLA) | 563 | 0.27 | 0.32 | 0.03 | 0.33 | 0.35 |
| C20:4n-6 | Arachidonic acid (AA) | 563 | 0.54 | 0.56 | 0.03 | 0.59 | 0.28 |
| C22:4n-6 | Adrenic acid | 561 | 0.21 | 0.26 | 0.03 | 0.23 | 0.20 |

Median, mean and variance showed the distribution of the proportion of each type of fatty acid in the subcutaneous adipose biopsy; N, sample size; The heritability ( $h^2$ ) was estimated by the ACE model, adjusted for age among 486 twins (210 MZ and 276 DZ) and 390 twins (214 DZ and 176 MZ) for adipose and serum fatty acid, respectively.

### The heritability of fatty acids is highly variable and tissue-specific

#### Adipose fatty acid

To understand how genetic variation contributes to adipose fatty acid contents, we utilized classic twin models to calculate the heritability of 18 individual fatty acids and the summary values of SFA, MUFA, and PUFA (Figure 1). We observed a broad range of heritability (0 to 59%, Table 1), with the highest heritability observed for arachidonic acid (AA). Overall, individual PUFAs showed consistent moderate to high heritability (h^2^ = 23 - 59%), with the exception of eicosapentaenoic acid (EPA) and docosahexaenoic acid (DHA) (h^2^ ∼ 0%). Conversely, only two of six SFAs (palmitic acid and stearic acid) were highly heritable (h^2^ = 30% and 51%, respectively), while the remaining SFAs and all three MUFAs had low heritability (h^2^ < 10%).

Next, we calculated the heritability of 41 product-to-precursor ratios of fatty acids. Once more, we observed a broad range of heritability (h^2^ = 0 - 54%) (Table S3). Fifteen product-to-precursor ratios were heritable (h^2^ ≥ 15%); and the mean heritability of the product-to-precursor ratios (h^2^ = 29%) was greater than the mean heritability of the individual level fatty acids described above (Table 1), suggesting genetic factors contribute relatively more to the enzymatic activity and conversion of fatty acids than their absolute levels.

#### Serum fatty acid

To compare the relative contribution of genetic regulation of fatty acids in different tissues, we investigated the heritability of 15 fatty acid levels in serum samples from a subset of the TwinsUK individuals (195 pairs of twins), which were profiled using the Metabolon platform on the same day as their adipose biopsy. Consistent with our adipose results, fatty acids in serum showed a range of heritabilities (h^2^ = 5 - 49%) (Table 1). While we saw similarity in the heritability estimates across serum and adipose tissue for some fatty acids (e.g. palmitic acid, docosapentaenoic acid, linoleic acid, dihomo-γ-linolenic acid and adrenic acid), others showed great discrepancies in heritability across tissues. For example, myristic acid was highly heritable (h^2^ = 44%) in serum yet was not heritable in adipose tissue (h^2^ = 0%), whereas both arachidonic acid and α-linolenic acid showed substantially higher heritability in adipose than serum (Table 1). Notably, the most common fatty acid in adipose, oleic acid, showed no heritability in adipose, but high heritability in serum (h^2^ = 40%). Overall, these results indicated substantial genetic regulation of fatty acids in both adipose tissue and serum, with some important tissue-specific effects. We saw no difference in heritability results when including BMI and/or cell type composition as covariates (Table S3 and Table S4).

### Identification of genetic variants associated with adipose fatty acids

In order to identify genetic loci associated with adipose fatty acid levels, we performed GWASs of 18 individual fatty acids, three fatty acid sums (SFA, MUFA, and PUFA) and 15 ratios of product-to-precursor fatty acids (h^2^ ≥ 0.15) (Figure 1). We identified ten independent genome-wide significant associated loci (*P_G_* < 5 × 10^−8^) across six fatty acids and seven fatty acid ratios (Table 2, Figure 2, Figure S2 and Figure S3), eight of which were novel discoveries, which do not overlap with any locus previously reported as genome-wide significant for fatty acids in adipose tissue. Two of ten associated loci (*SCD locus* - lead variant rs603424, and 3p25.2 *locus* - lead variant rs6768977) were significant at a study-wide multiple testing corrected threshold accounting for the number of independent fatty acids (*P_B_* < 2.8 ×10^−9^), among which 3p25.2 *locus* (rs6768977) was a novel finding. The *SCD locus* (rs603424) was associated with SFA, palmitic acid, stearic acid, palmitoleic acid, the ratio of palmitoleic acid/palmitic acid and the ratio of oleic acid/stearic acid. The lead variant explained 7.0 - 11.2% of the variance in SFA and MUFA. Variants at the *FADS* cluster have been widely reported to regulate the desaturation of PUFAs, in this study we identified associations to arachidonic acid and the ratio of arachidonic acid/dihomo-γ-linolenic acid. The lead variants at the *FADS locus*, rs97384 and rs174544 were located in the protein-coding gene *FADS2*, close to *FADS1*/*FEN1*/*TMEM258*, and explained 6.5% and 7.9% of the variance. By conditioning on the lead variant at the *FADS locus* (rs97384), we found four secondary variants at the *FADS locus* where the lead variant was rs61898565 (*P* = 6.3 ×10^−5^).

**Figure 2.**
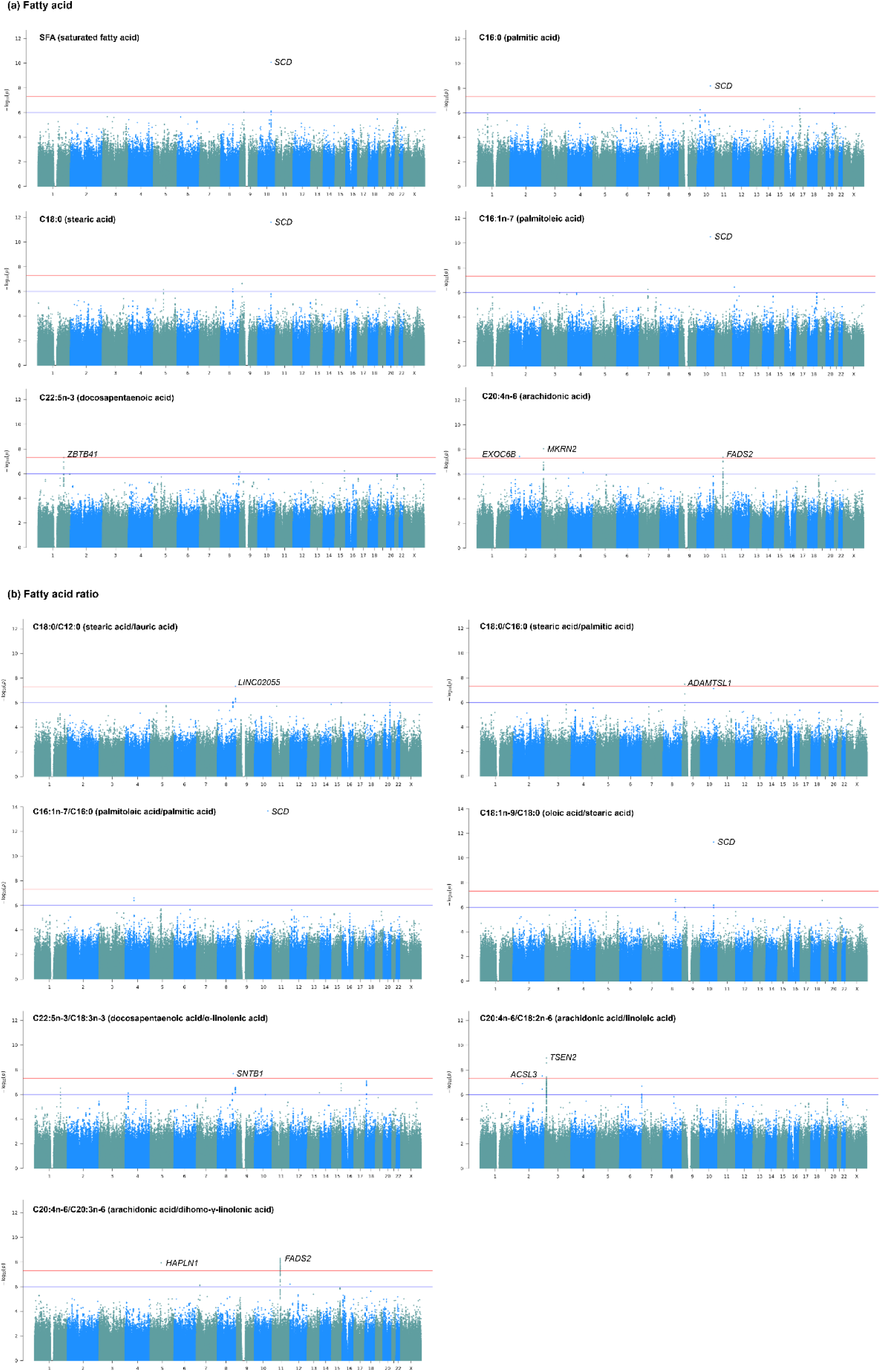
Manhattan plots of GWASs on fatty acids in adipose tissue. X-axis indicates chromosome 1 to 22 and chromosome X; Y-axis indicates minus log10-transformed GWAS *P* value; the red horizontal line demarcates the genome-wide significance threshold of *P_G_* = 5 x 10^-8^, the genome-wide significant loci are marked and annotated to the nearest genes, the blue line indicates suggestive significance threshold of *P_s_* = 10^-6^.

**Table 2.** Genome-wide significant loci of fatty acids in adipose tissue.

| Adipose fatty acid | Variant | Chr | Pos | Nearest genes | Effect allele | Non-effect allele | EAF | Effect | SE | P | Variance explained by the variant (%) |
| --- | --- | --- | --- | --- | --- | --- | --- | --- | --- | --- | --- |
| **Fatty acid** |  |  |  |  |  |  |  |  |  |  |  |
| SFA (saturated fatty acid) | rs603424 | 10 | 100,315,722 | *PKD2L1/SCD* | A | G | 0.176 | 0.554 | 0.084 | 8.5E-11<sup>a</sup> | 9.0 |
| C16:0 (palmitic acid) | rs603424 | 10 | 100,315,722 | *PKD2L1/SCD* | A | G | 0.175 | 0.490 | 0.083 | 6.6E-09 | 7.0 |
| C18:0 (stearic acid) | rs603424 | 10 | 100,315,722 | *PKD2L1/SCD* | A | G | 0.175 | 0.596 | 0.083 | 2.4E-12<sup>a</sup> | 10.1 |
| C16:1n-7 (palmitoleic acid) | rs603424 | 10 | 100,315,722 | *PKD2L1/SCD* | A | G | 0.175 | -0.542 | 0.080 | 3.1E-11<sup>a</sup> | 8.6 |
| C22:5n-3 (docosapentaenoic acid) | rs114713999 | 1 | 197,160,467 | *ZBTB41/ASPM* | A | G | 0.012 | 1.521 | 0.275 | 4.8E-08<sup>b</sup> | 5.5 |
| C20:4n-6 (arachidonic acid) | rs17559539 | 2 | 72,348,460 | *EXOC6B/CYP26B1* | C | A | 0.044 | -0.955 | 0.171 | 3.7E-08<sup>b</sup> | 7.7 |
| C20:4n-6 (arachidonic acid) | rs1542848 | 3 | 12,576,497 | *MKRN2/RAF1/TSEN2/PPARG* | A | G | 0.375 | -0.379 | 0.065 | 8.7E-09<sup>b</sup> | 6.8 |
| C20:4n-6 (arachidonic acid) | rs97384 | 11 | 61,856,709 | *FADS2/FADS1/FADS3/FEN1/TMEM258* | C | T | 0.621 | 0.370 | 0.067 | 4.3E-08 | 6.5 |
| **Fatty acid ratio** |  |  |  |  |  |  |  |  |  |  |  |
| C18:0/C12:0 (stearic acid/lauric acid) | rs4460408 | 8 | 137,045,124 | *LINC02055/RNU6-144P* | A | C | 0.780 | 0.439 | 0.079 | 4.6E-08<sup>b</sup> | 6.6 |
| C18:0/C16:0 (stearic acid/palmitic acid) | rs776749 | 9 | 18,642,756 | *ADAMTSL1/MIR3152* | A | G | 0.535 | 0.372 | 0.066 | 3.3E-08<sup>b</sup> | 6.8 |
| C16:1n-7/C16:0 (palmitoleic acid/palmitic acid) | rs603424 | 10 | 100,315,722 | *PKD2L1/SCD* | A | G | 0.175 | -0.624 | 0.079 | 2.2E-14<sup>a</sup> | 11.2 |
| C18:1n-9/C18:0 (oleic acid/stearic acid) | rs603424 | 10 | 100,315,722 | *PKD2L1/SCD* | A | G | 0.175 | -0.587 | 0.083 | 5.1E-12<sup>a</sup> | 9.8 |
| C22:5n-3/C18:3n-3 (docosapentaenoic acid/α-linolenic acid) | rs79670584 | 8 | 120,585,786 | *SNTB1/MTBP/MRPL13/COL14A1* | A | T | 0.019 | 1.410 | 0.248 | 2.1E-08<sup>b</sup> | 7.4 |
| C20:4n-6/C18:2n-6 (arachidonic acid/linoleic acid) | rs795891 | 2 | 222,881,016 | *ACSL3/MOGAT1* | G | A | 0.820 | 0.480 | 0.085 | 3.1E-08<sup>b</sup> | 6.9 |
| C20:4n-6/C18:2n-6 (arachidonic acid/linoleic acid) | rs6768977 | 3 | 12,466,670 | *TSEN2/PPARG/MKRN2/RAF1* | G | A | 0.494 | -0.379 | 0.061 | 1.1E-09<sup>a,b</sup> | 7.2 |
| C20:4n-6/C20:3n-6 (arachidonic acid/dihomo-γ-linolenic acid) | rs79680695 | 5 | 83,784,409 | *HAPLN1/EDIL3* | G | C | 0.050 | 0.854 | 0.147 | 1.2E-08<sup>b</sup> | 7.0 |
| C20:4n-6/C20:3n-6 (arachidonic acid/dihomo-γ-linolenic acid) | rs174544 | 11 | 61,800,281 | *FADS2/FADS1/FEN1/TMEM258/FADS3* | A | C | 0.307 | -0.429 | 0.072 | 4.8E-09 | 7.9 |

Chr, chromosome; Pos, position (GRCh38/hg38); EAF, effect allele frequency; Effect, beta estimate of effect allele; SE, standard error; Variance explained, the variance explained by the lead variant. For each locus, only the lead variant was reported here. <sup>a</sup> Bonferroni-corrected genome-wide significance with $P_B < 2.8 \times 10^{-9}$ ; <sup>b</sup> Novel locus.

Six out of eight novel loci were associated with polyunsaturated fatty acids. The 3p25.2 *locus* (rs1542848 and rs6768977) was associated with arachidonic acid and the ratio of arachidonic acid/linoleic acid. The *CYP26B1 locus* (rs795891) was associated with arachidonic acid. A total of 266 SNPs reached the suggestive threshold (*P_S_* < 10^−6^) (Table S5), suggesting higher powered GWAS may robustly identify further genetic associations.

In order to test the robustness of the approach, we have also performed sensitivity analyses by re-running all heritability estimates and GWASs including the potential confounders, BMI and/or cell types, as covariates. Overall, we see similar results across the analyses, with slight fluctuation of *P* values for some fatty acids (Table S3, Table S4 and Table S6). Given the consistency in the results, all further analyses in the main text reference the GWAS results that do not adjust for BMI or cell type composition.

### Replication and tissue specificity

We sought to replicate the genome-wide significant GWAS loci in both adipose tissue and serum/plasma datasets. The only published GWAS of fatty acids in adipose tissue investigated the association between estimates of adipose desaturase activity for D5D, D6D and SCD as measured by fatty acid product to precursor ratios in buttock subcutaneous adipose tissue obtained from 783 men from ULSAM study ^13^. This study reported genome-wide significant associations of D5D (C20:4n-6/C20:3n-6 ratio) to the *FADS locus* and SCD (C16:1/C16:0 ratio) to the *SCD locus*, which are consistent with our results at these loci. A second study investigating candidate SNPs at the *FADS locus* with adipose tissue fatty acid levels in 173 European individuals from the DiOGenes also reported an association with D5D activity (C20:4n-6/C18:2n-6 ratio) ^12^. However, the full summary statistics of the previous adipose fatty acid studies could not be made available, so we could not investigate further replication in adipose tissue. Therefore, we sought to replicate the GWAS results in serum metabolite datasets.

We first performed GWASs of matched serum metabolite dataset in TwinsUK where blood samples were collected at the same time as adipose tissue biopsies within the same group of participants (N = 448). We replicated the associations between *SCD locus* (rs603424) and *FADS locus* (rs97384) and three fatty acids (Table 3). We then sought to replicate the GWAS results in four larger cohorts including a combined TwinsUK and KORA cohort (N = 7,824) ^24^, a combined EPIC-Norfolk and INTERVAL cohort (N = 19,994) ^25^, and CLSA cohort (N = 8,299) ^26^. We replicated the association between *SCD locus* (rs603424) and stearic acid and palmitoleic acid, and the associations between *FADS locus* (rs97384 and its LD proxies, rs174548 and rs102275) and arachidonic acid level. Together, these results suggest the majority of testable loci are replicated which show shared metabolic pathways in both serum and adipose tissue.

**Table 3.**
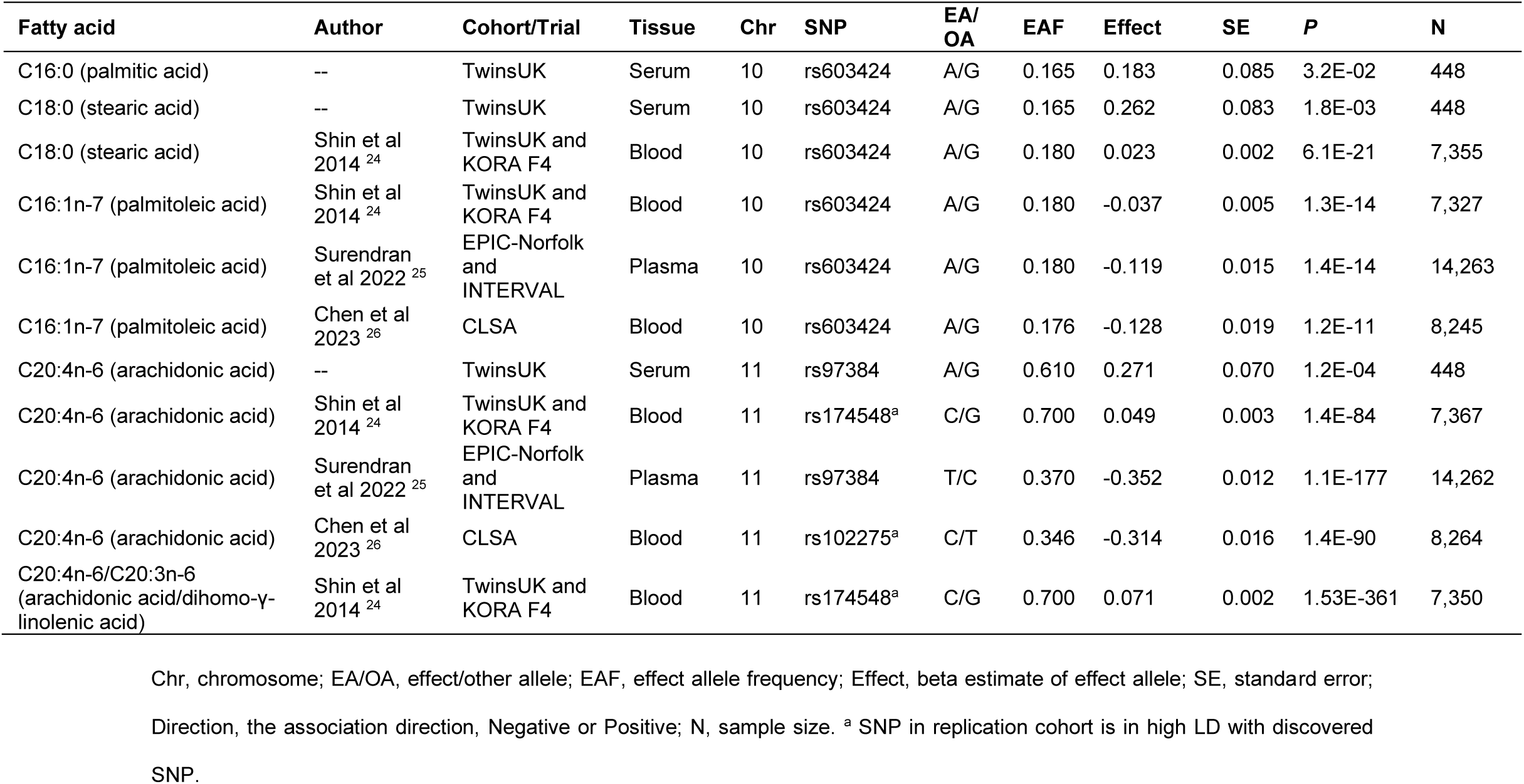
Replicated variant-fatty acid associations in other cohorts.

| Fatty acid | Author | Cohort/Trial | Tissue | Chr | SNP | EA/OA | EAF | Effect | SE | P | N |
| --- | --- | --- | --- | --- | --- | --- | --- | --- | --- | --- | --- |
| C16:0 (palmitic acid) | -- | TwinsUK | Serum | 10 | rs603424 | A/G | 0.165 | 0.183 | 0.085 | 3.2E-02 | 448 |
| C18:0 (stearic acid) | -- | TwinsUK | Serum | 10 | rs603424 | A/G | 0.165 | 0.262 | 0.083 | 1.8E-03 | 448 |
| C18:0 (stearic acid) | Shin et al 2014<sup>24</sup> | TwinsUK and KORA F4 | Blood | 10 | rs603424 | A/G | 0.180 | 0.023 | 0.002 | 6.1E-21 | 7,355 |
| C16:1n-7 (palmitoleic acid) | Shin et al 2014<sup>24</sup> | TwinsUK and KORA F4 | Blood | 10 | rs603424 | A/G | 0.180 | -0.037 | 0.005 | 1.3E-14 | 7,327 |
| C16:1n-7 (palmitoleic acid) | Surendran et al 2022<sup>25</sup> | EPIC-Norfolk and INTERVAL | Plasma | 10 | rs603424 | A/G | 0.180 | -0.119 | 0.015 | 1.4E-14 | 14,263 |
| C16:1n-7 (palmitoleic acid) | Chen et al 2023<sup>26</sup> | CLSA | Blood | 10 | rs603424 | A/G | 0.176 | -0.128 | 0.019 | 1.2E-11 | 8,245 |
| C20:4n-6 (arachidonic acid) | -- | TwinsUK | Serum | 11 | rs97384 | A/G | 0.610 | 0.271 | 0.070 | 1.2E-04 | 448 |
| C20:4n-6 (arachidonic acid) | Shin et al 2014<sup>24</sup> | TwinsUK and KORA F4 | Blood | 11 | rs174548<sup>a</sup> | C/G | 0.700 | 0.049 | 0.003 | 1.4E-84 | 7,367 |
| C20:4n-6 (arachidonic acid) | Surendran et al 2022<sup>25</sup> | EPIC-Norfolk and INTERVAL | Plasma | 11 | rs97384 | T/C | 0.370 | -0.352 | 0.012 | 1.1E-177 | 14,262 |
| C20:4n-6 (arachidonic acid) | Chen et al 2023<sup>26</sup> | CLSA | Blood | 11 | rs102275<sup>a</sup> | C/T | 0.346 | -0.314 | 0.016 | 1.4E-90 | 8,264 |
| C20:4n-6/C20:3n-6 (arachidonic acid/dihomo-γ-linolenic acid) | Shin et al 2014<sup>24</sup> | TwinsUK and KORA F4 | Blood | 11 | rs174548<sup>a</sup> | C/G | 0.700 | 0.071 | 0.002 | 1.53E-361 | 7,350 |

Chr, chromosome; EA/OA, effect/other allele; EAF, effect allele frequency; Effect, beta estimate of effect allele; SE, standard error; Direction, the association direction, Negative or Positive; N, sample size. <sup>a</sup> SNP in replication cohort is in high LD with discovered SNP.

### GWAS signals are colocalized with eQTL and meQTL signals and highlight tissue specificity

Although GWASs can identify genetic variants associated with phenotypes or diseases, the molecular mechanisms underlying these associations often remain unclear. Colocalization of GWAS and molecular trait genetic associations (such as eQTLs or meQTLs) can assess whether the same variants influence both a molecular trait and adipose fatty acids, potentially highlighting causal mechanisms. We integrated the GWAS signals with three molecular datasets: 1) adipose eQTLs generated from the same TwinsUK adipose biopsies as the fatty acid quantifications, 2) adipose methylation QTLs (meQTLs) generated from the same TwinsUK biopsies and, and 3) eQTLs from the multi-tissue GTEx study. In brief, we found three loci at *SCD*, *FADS* and 3p25.2 influenced both molecular traits and adipose fatty acids.

#### TwinsUK eQTL-GWAS colocalization

We integrated the concurrent adipose tissue eQTL data and our adipose fatty acid GWAS data to perform colocalization analysis to prioritize candidate causal genes underlying the GWAS loci (Figure 1). Overall, we identified 11 colocalizations cases where the gene expression and GWAS trait are driven by the same genetic variant, rather than by variants in linkage disequilibrium (with PP.H4 > 0.75), spreading across two GWAS loci and regulating the expression of four genes (Table 4 and Figure 3). GWAS-eQTL colocalizations were: The GWAS signal of SFA and MUFA (*SCD locus*) colocalized with the primary eQTL signal of *SCD*, and GWAS signals of n-6 PUFA (*FADS locus*) colocalized with primary eQTLs signals (or signals in high LD with eQTL signals) of *FEN1*, *FADS1* and *TMEM258*. All colocalizations passed sensitivity analyses (Figure S4). As the *SCD locus* (rs603424) is the single independent GWAS signal (no other SNPs in high LD nearby) of SFA and MUFA and this signal explained around 10% of variance, we hypothesized this partially due to the high heritability of *SCD* expression (h^2^ = 42%). The secondary eQTL signal of *MKRN2* colocalized with GWAS signal of arachidonic acid. Together, these results demonstrate evidence of the functional effects of the GWAS signals that are mediated through the expression of *SCD*, *FADS1*, *FEN1* and *TMEM258*.

**Figure 3.**
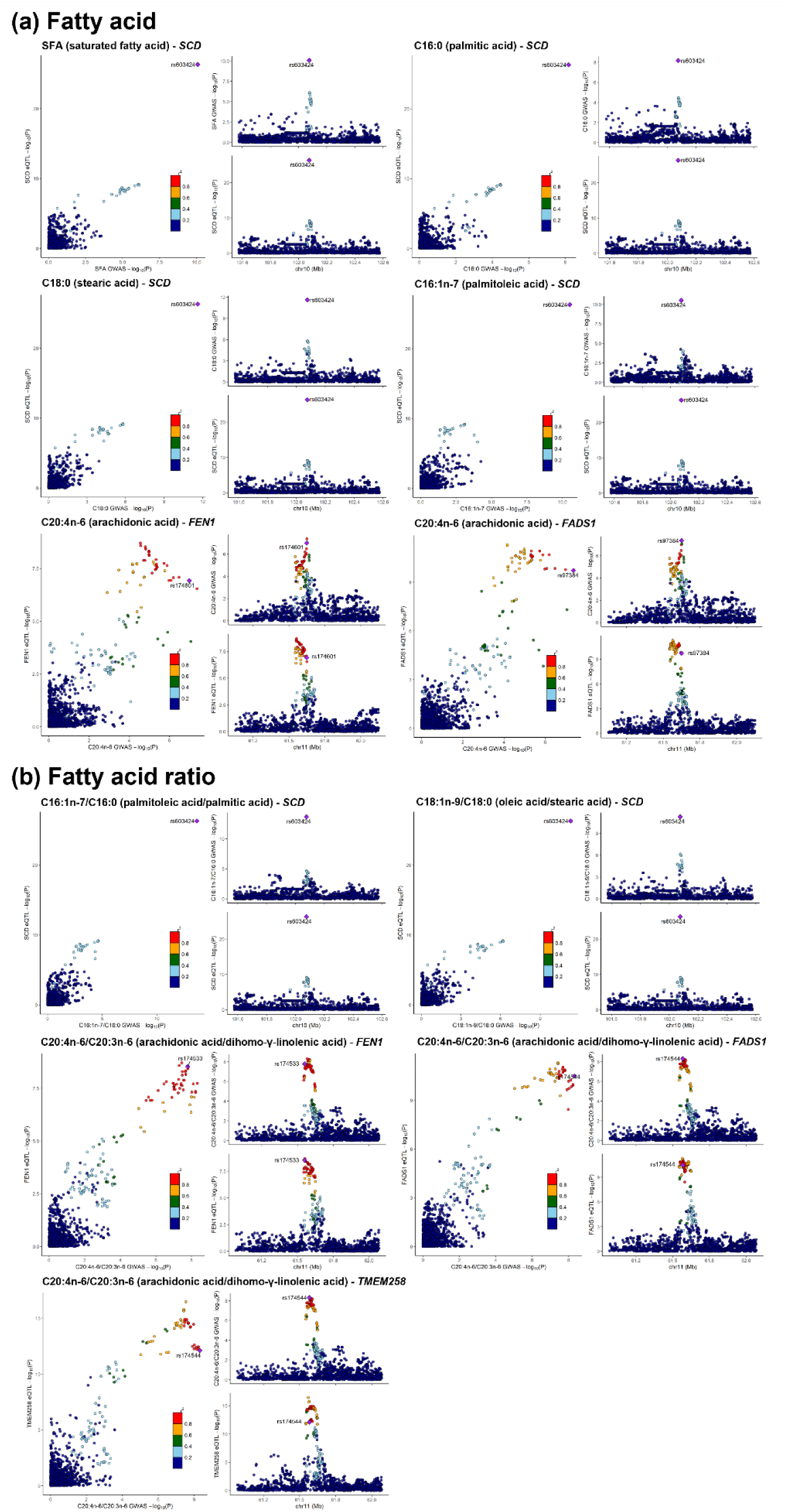
Colocalization of GWAS and eQTL signals. The labeled SNP is the lead SNP for both GWAS and eQTL studies, and other SNPs are colored according to their LD r^2^ with the lead SNP.

**Table 4.** Colocalization of GWAS and eQTL signals.

| Adipose fatty acid | Chr | Lead GWAS SNP | Ensembl ID | Gene symbol | Lead eQTL SNP | LD $r^2$ | PP.H4 | Colocalized SNP |
| --- | --- | --- | --- | --- | --- | --- | --- | --- |
| **Fatty acid** |  |  |  |  |  |  |  |  |
| SFA (saturated fatty acid) | 10 | rs603424 | ENSG00000099194 | *SCD* | rs603424 | 1.00 | 1.00 | rs603424 |
| C16:0 (palmitic acid) | 10 | rs603424 | ENSG00000099194 | *SCD* | rs603424 | 1.00 | 1.00 | rs603424 |
| C18:0 (stearic acid) | 10 | rs603424 | ENSG00000099194 | *SCD* | rs603424 | 1.00 | 1.00 | rs603424 |
| C16:1n-7 (palmitoleic acid) | 10 | rs603424 | ENSG00000099194 | *SCD* | rs603424 | 1.00 | 1.00 | rs603424 |
| C20:4n-6 (arachidonic acid) | 11 | rs97384 | ENSG00000168496 | *FEN1* | rs174536 | 0.76 | 0.79 | rs174601 |
| C20:4n-6 (arachidonic acid) | 11 | rs97384 | ENSG00000149485 | *FADS1* | rs4246215 | 0.68 | 0.86 | rs97384 |
| C20:4n-6 (arachidonic acid) | 3 | rs1542848 | ENSG00000075975 | *MKRN2*<sup>a</sup> | rs299628 | 0.63 | 0.76 | rs1542848 |
| **Fatty acid ratio** |  |  |  |  |  |  |  |  |
| C16:1n-7/C16:0<br>(palmitoleic acid/palmitic acid) | 10 | rs603424 | ENSG00000099194 | *SCD* | rs603424 | 1.00 | 1.00 | rs603424 |
| C18:1n-9/C18:0<br>(oleic acid/stearic acid) | 10 | rs603424 | ENSG00000099194 | *SCD* | rs603424 | 1.00 | 1.00 | rs603424 |
| C20:4n-6/C20:3n-6<br>(arachidonic acid/dihomo-γ-<br>linolenic acid) | 11 | rs174544 | ENSG00000168496 | *FEN1* | rs174536 | 0.81 | 0.96 | rs174533 |
| C20:4n-6/C20:3n-6<br>(arachidonic acid/dihomo-γ-<br>linolenic acid) | 11 | rs174544 | ENSG00000134825 | *TMEM258* | rs174538 | 0.67 | 0.95 | rs174544 |
| C20:4n-6/C20:3n-6<br>(arachidonic acid/dihomo-γ-<br>linolenic acid) | 11 | rs174544 | ENSG00000149485 | *FADS1* | rs4246215 | 0.75 | 0.95 | rs174544 |

LD $r^2$ , linkage disequilibrium between GWAS and eQTL lead SNPs; PP.H4, posterior probability of hypothesis 4, both traits are associated and share a single causal variant; Colocalized SNP, the lead colocalized SNP for both GWAS and eQTL studies; <sup>a</sup> Secondary signal of the gene *MKRN2*.

#### GTEx QTL-GWAS colocalization across multiple tissues

To investigate overlap with genetic regulation of gene expression across multiple tissues, we performed colocalization between the adipose fatty acids GWAS and GTEx eQTL and sQTL. We found association between rs603424 and *SCD* expression was specific to adipose tissue; present only in GTEx visceral and subcutaneous adipose tissue and a weaker association in breast mammary tissue (which contains adipocytes). In contrast, the eQTLs at the *FADS* cluster were shared across multiple tissues, including adipose subcutaneous, adipose visceral omentum, pancreas, lung, artery tibial, colon transverse, verve tibial, esophagus muscularis, esophagus mucosa, and esophagus gastroesophageal junction. This analysis also revealed that eQTLs at *RAF1* and *TSEN2* genes had shared effects across nerve tibial, pituitary, adipose visceral omentum, adrenal gland, colon sigmoid, cells EBV-transformed lymphocytes and artery tibial (Table S7 and Table S8).

#### TwinsUK meQTL-GWAS colocalization

To identify potential methylation of CpG sites mediating the associations between genetic variants and fatty acid levels, we integrated the GWAS loci with adipose meQTLs generated from the same TwinsUK individuals (Figure 1). Using a Bayesian colocalization approach, we identified 71 meQTL-GWAS colocalizations across nine fatty acids and 24 CpGs (Table S9, Figure S5, and Figure S6). We identified *SCD locus* influencing both the DNA methylation level of eight CpGs near *SCD* and SFA and MUFA levels in adipose tissue. The 3p25.2 *locus* influenced both the DNA methylation level of seven CpGs near *MKRN2/TSEN2* and arachidonic acid and arachidonic acid/linoleic acid ratio. Colocalizations were identified between CpGs near *FADS1/FADS2* and the primary GWAS signals at this locus, but no colocalizations were identified for the secondary GWAS signal.

### Adipose DNA methylation, gene expression and fatty acid cross genotypes

To further explore genetic regulation of fatty acid levels and ratios at the loci with evidence of colocalizations, we tested the associations between concurrent DNA methylation, gene expression and fatty acid in adipose tissue and plotted concurrent measures across genotypes. Adipose fatty acids were associated with the expression levels of *SCD*, *FEN1*, *FADS1* and *TMEM258* (Table S10) and DNA methylation levels of 24 CpG sites near *SCD/FADS1/TSEN2* genes (Table S11). For example, with the increase in alternative allele frequency of rs603424 (A), the methylation level of cg01270221 (located within the gene body) increased, the expression level of *SCD* gene decreased, and the ratio of palmitoleic acid to palmitic acid decreased (Figure 4 and Figure S7). The allele of rs174544_A was associated with lower levels of cg25448062 (located within the gene body), *FADS1* expression and the ratio of arachidonic acid/dihomo-γ-linolenic acid in adipose tissue.

**Figure 4.**
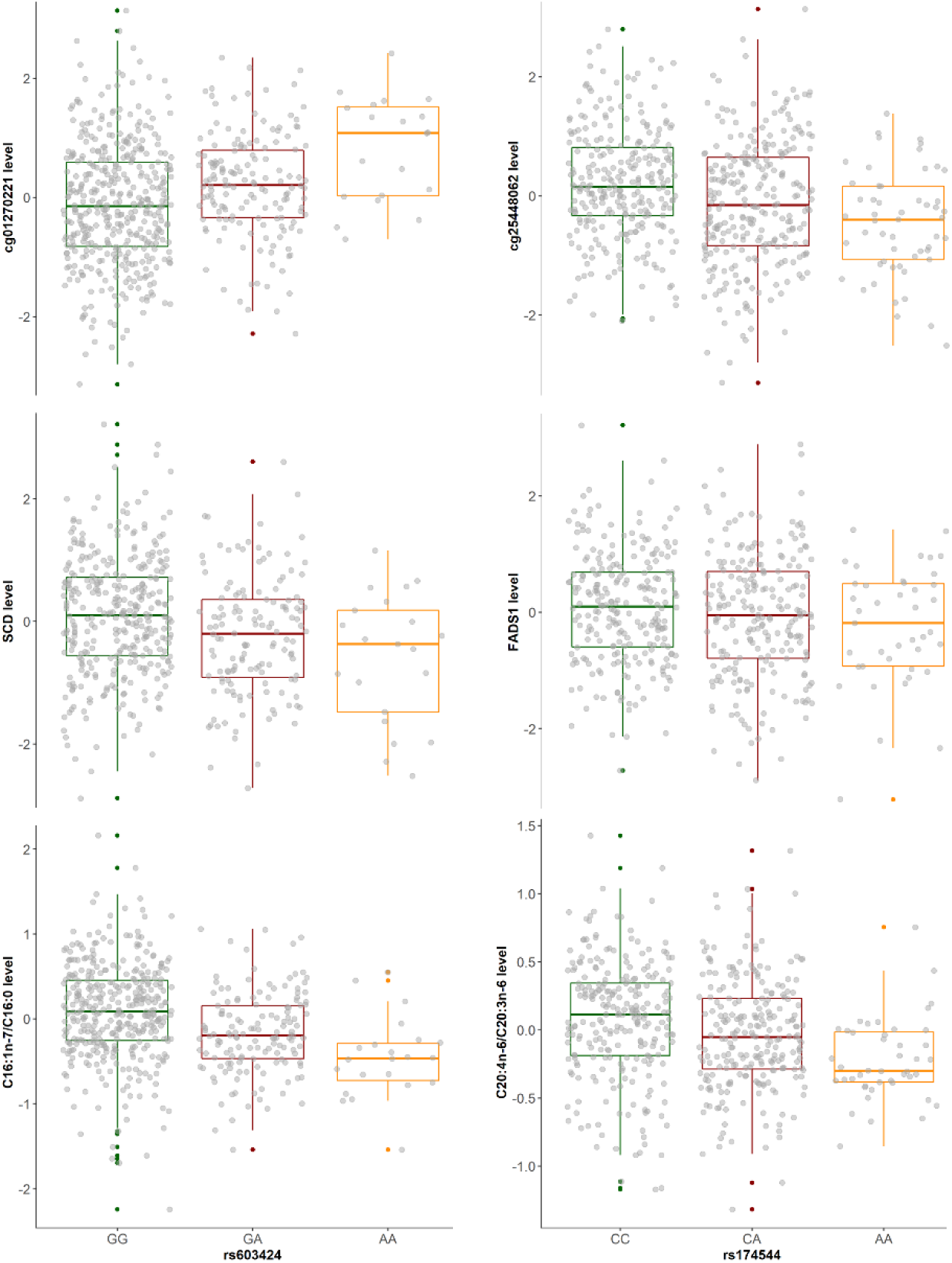
Box plots of levels of DNA methylation, gene expression and fatty acids on genotypes for colocalized GWAS-eQTL or meQTL signals. The left plot shows the levels of cg01270221, *SCD* gene and C16:1n-7/C16:0 (palmitoleic acid/palmitic acid) by the SNP rs603424 in adipose tissue; the right plot shows the levels of cg25448062, *FADS1* gene and C20:4n-6/C20:3n-6 (arachidonic acid/dihomo-γ-linolenic acid) by the SNP rs174544 in adipose tissue. We take the most significant fatty acid or CpG associated with the same SNP as the examples here. We regress out age and family relatedness for fatty acid; regress out technical covariates, age, BMI, and family relatedness for gene expression; regress out technical covariates, age, predicted smoking, family relatedness, genetic PCs, and non-genetic DNA methylation PCs for DNA methylation.

### Fatty acids and kidney traits shared causal variants

In this study, we demonstrated a study-wide significant association between the genetic *locus* 3p25.2 (near *PPARG*) and adipose tissue levels of arachidonic acid and arachidonic acid/linoleic acid ratio. A recent review has highlighted the implication of PPARγ in critical conditions such as pulmonary arterial hypertension and kidney failure ^53^. eGFR, creatinine and cystatin C are most widely used renal biomarkers ^54^ and available in TwinsUK cohorts. To identify whether the genetic *locus* 3p25.2 influence both adipose fatty acids and kidney traits, we colocalized Sinnott-Armstrong et al. GWAS summary statistics of three kidney traits (eGFR, creatinine and cystatin C) ^43^ with TwinsUK adipose fatty acid GWAS results. We found that eGFR, serum creatinine and cystatin C shared the single causal variants (rs9825233/rs904464) with arachidonic acid level in adipose tissue, with the posterior probabilities ranging from 0.91 to 0.98. The GWAS signal rs904464 had an effect on both cystatin C level and arachidonic acid/linoleic acid ratio with the posterior probability of 0.99. These suggest that the same genetic *locus* 3p25.2 influences both adipose n-6 PUFAs and kidney health (Figure 5 and Figure S8).

**Figure 5.**
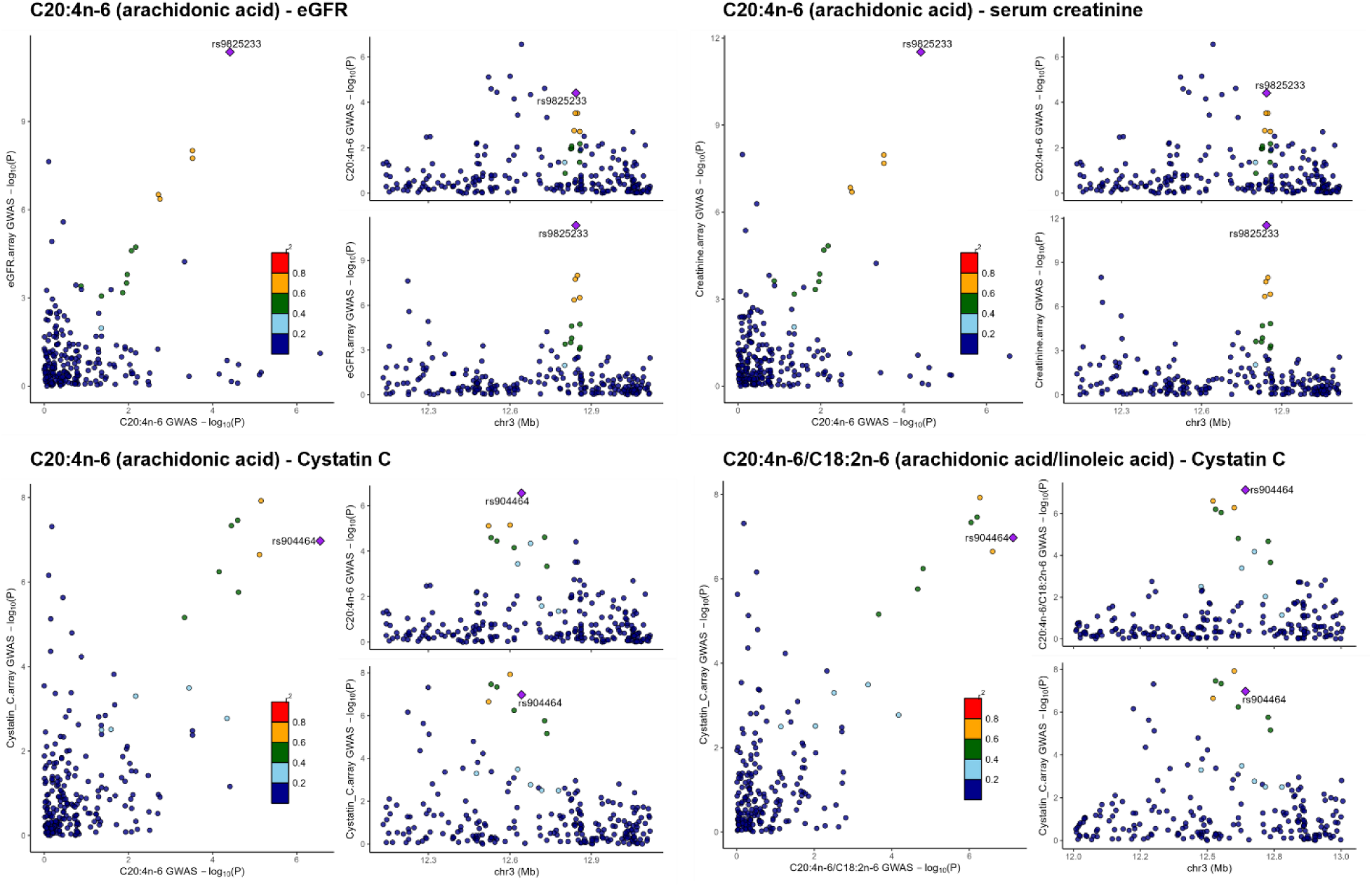
Colocalization of TwinsUK fatty acid GWAS signals and published kidney trait GWAS signals. The labeled variant is the lead variant for both GWAS studies, and other variants are colored according to their LD r^2^ with the lead variant.

### Genetic regulation of fatty acid links with biomedical phenotypes

To explore the links between genetic regulation of fatty acids and other biomedical phenotypes, we conducted phenome-wide association studies (PheWASs) of the 12 GWAS lead variants using NIH Accelerating Medicines Partnership (AMP) and GWASATLAS databases (Figure 1). We determined that 95 traits and 128 traits in AMP and GWASATLAS databases, respectively, were associated with adipose fatty acid-associated variants (*P* < 10^-5^) (Table S12). Saturated and monounsaturated fatty acids-associated variant rs603424 at the *SCD locus* was also associated with mean platelet volume, body fat percentage, waist-hip ratio, bone mineral density, FEV1/FVC ratio, and cholesterol (Figure S9). The variant rs174544 at *FADS locus* associated with arachidonic acid/dihomo-γ-linolenic acid ratio in this study was reported to be associated with total cholesterol, HDL and LDL, triglycerides, alkaline phosphatase, mean platelet volume, red and white blood cell count, hemoglobin concentration, fasting glucose, HbA1c, T2D, pulse rate, plasma C-reactive protein and sex hormone binding globulin levels (Figure S10). These results suggest that SNPs regulating adipose fatty acids are also involved in cardio-metabolic traits and therefore could be implicated in disease risk.

To further investigate how regulation of fatty acids may correlate with the heritable risks of developing diseases, we utilized polygenic scores (PGSs) for adiposity traits (visceral, abdominal subcutaneous, and gluteofemoral adipose tissue volumes and waist-to-hip ratio), BMI, lipid biomarkers (HDL, LDL, total triglycerides and total cholesterol) and chronic diseases (type 1 and 2 diabetes, coronary artery disease and hypertension) and fatty acids in adipose tissue (Figure 1 and Table S1). The PGSs were first validated by the corresponding trait measures in the full population of TwinsUK (Table S2). We found significant associations between fatty acid levels in adipose tissue and PGSs for BMI, gluteofemoral adipose tissue volumes, HDL and total triglycerides (Figure 6). More specifically, we observed a negative association of PGS BMI to three saturated fatty acids and a positive association of PGS BMI to three unsaturated fatty acids. The PGS for gluteofemoral adipose tissue volumes showed a positive association with palmitoleic acid level. Interestingly, individuals with high levels of lauric acid showed a pattern of elevated PGS HDL and reduced PGS total triglycerides (all adjusted *P* < 0.05, Figure 6 and Table S13). In conclusion, polygenic scores of cardio-metabolic traits were associated with adipose tissue fatty acid levels, suggesting genetic risk for these traits also influence fatty acid composition in this key metabolic tissue.

**Figure 6.**
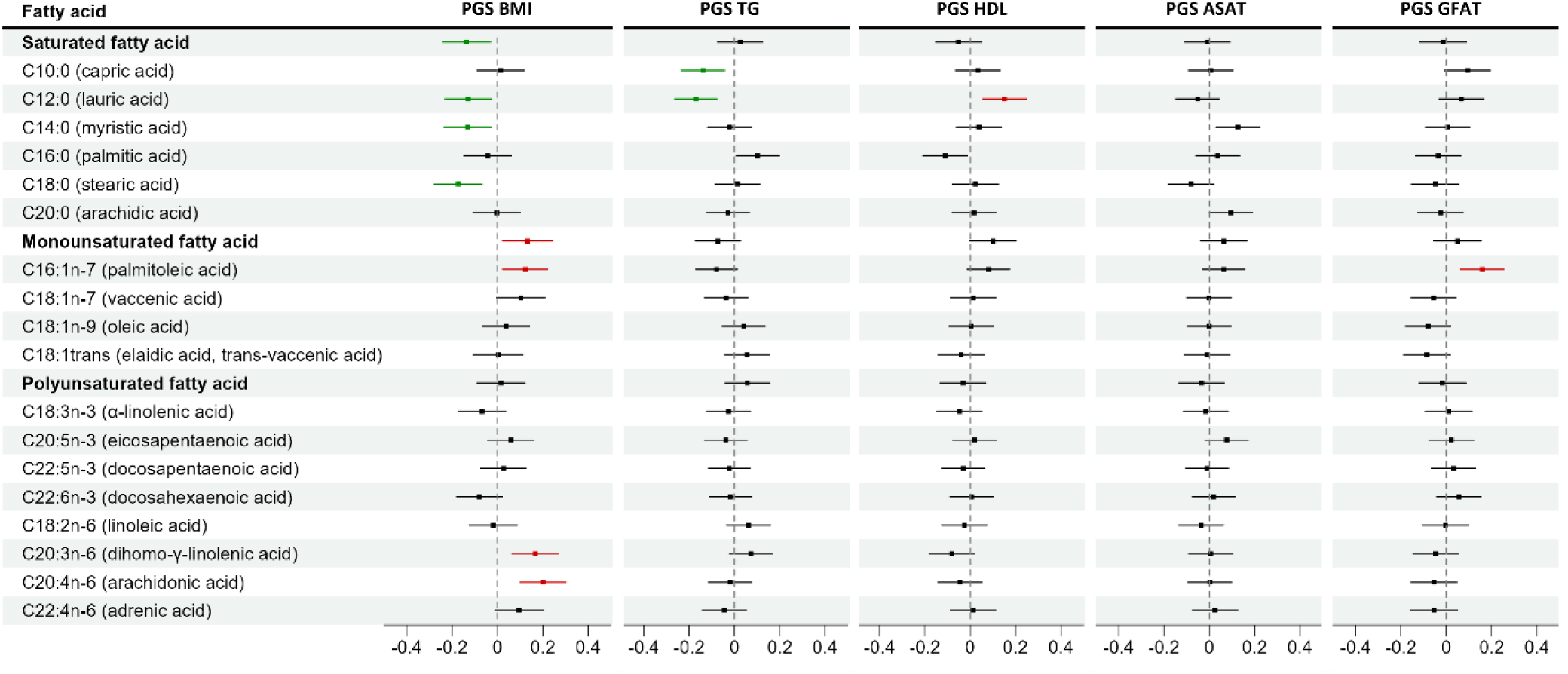
Forest plot of association between polygenic scores and fatty acids in adipose tissue. The dots display effect sizes and horizontal lines show their confidence intervals. Significant associations are marked red (positive) and green (negative). BMI, body mass index; TG, total triglycerides; HDL, high-density lipoproteins cholesterol; ASAT, abdominal subcutaneous adipose tissue volumes; GFAT, gluteofemoral adipose tissue volumes.

## Discussion

In summary, we characterized the heritability of adipose fatty acids and identified ten GWAS loci associated with fatty acid levels or fatty acid product to precursor ratios in adipose tissue. Through multi-omics colocalization analyses, we prioritized 24 CpGs and five genes mediating the genetic regulation of fatty acids in adipose tissue. Further, using phenome-wide association analyses and polygenic scores, we linked genetic regulation of adipose fatty acids with biomedical phenotypes.

Fatty acids are important biological molecules, which act as the bedrock for almost all lipid structures and can represent a snapshot of metabolic state. Studies have revealed that circulating fatty acids are associated with T2D, CVD, metabolic syndrome, cognitive function, Alzheimer’s disease, and inflammatory biomarkers ^26,55–57^, but the health impact of fatty acids shows controversial directions. For example, a positive relationship between circulating saturated fatty acids (myristic, palmitic, and stearic acids) and incident T2D and CVD have been reported, but eicosapentaenoic acid and docosahexaenoic acid were protective for CVD ^58^. Individual fatty acid contents and fatty acid conversions may be tissue-specific, reflecting different lipoprotein classes and tissue specific modifications. Adipose tissue, which consists almost entirely of triglycerides, is the largest store of fatty acids but also has an endocrine role. However, the regulatory network of fatty acid mechanism in subcutaneous adipose tissue remains unclear.

We revealed a broad range of heritability for individual fatty acids and the ratios of fatty acids, indicating substantial genetic regulation of some adipose fatty acids. The heritability of serum fatty acids ^7^ is in line with TwinsUK large-scale study but the heritability of serum oleic acid, eicosapentaenoic acid and docosahexaenoic acid levels were greater than those in adipose tissue. Tissue-specific fatty acid heritability may be due to enzymatic activity differences across tissues. The activities of D5D and D6D converting polyunsaturated fatty acids are higher in serum than in adipose tissue ^52,59^. Some saturated fatty acids were not heritable in adipose tissue, because they are converted mainly to palmitic acid or primarily obtained from dietary intake, such as coconut oil, palm kernel oil, or bovine milk ^60^.

SNPs at *SCD* and the *FADS* loci were associated with both adipose fatty acids and circulating fatty acids. It is well documented that *FADS* gene cluster polymorphisms regulate polyunsaturated fatty acids synthesis by altering the activity of fatty acid desaturases (D5D and D6D) and tissue availability ^61^; similarly, *SCD* gene polymorphisms regulate the rate limiting enzyme catalyzing monounsaturated fatty acids from saturated fatty acids ^13^. These two loci were replicated in independent cohorts and the gene expression levels of *FADS1* and *SCD* mediated the fatty acid conversion in adipose tissue. When we investigated the genetic regulation of expressions of these genes across multiple tissues, we found that *SCD* is adipose tissue-specific but *FADS1/FADS2* is tissue-shared. In line with previous studies ^62–65^, we found the SNPs at *FADS loci* and *SCD* are associated with T2D and cardio-metabolic markers on PheWAS platforms. ^66^Another key set of enzymes in fatty acid regulation are the elongases including the *ELOVL* gene family. Studies in serum have identified associations between SNPs near *ELOVL2* and polyunsaturated fatty acids ^67^. However, we did not identify any association near *ELOVL* genes and polyunsaturated fatty acids with genome-wide significance in adipose tissue.

The challenge of post-GWAS analyses is to elucidate the underlying causal genes and mechanisms involved at identified loci. The most common strategy for identifying the molecular mechanisms at GWAS loci is integration or colocalization with gene expression or DNA methylation quantitative trait loci datasets, which can identify the causal genes, epigenetic elements and/or tissue of action. This strategy has been successfully applied to previous loci such as *FADS locus*, where *FADS1* and *FADS2* expression and DNA methylation of CpG sites in *FADS1* and *FADS2* are associated with fatty acid desaturation in serum and adipose tissue ^12,68,69^. Further, some studies have implied the mediator roles of CpGs in the association between gene expression and serum fatty acid regulation ^70–72^. For example, a study conducted in a Chinese cohort has demonstrated that DNA methylation levels of four CpG sites mediated the effect of rs174570 on *FADS1* gene expression, while only DNA methylation of one CpG site mediated the effect of rs174570 on *FADS2* gene expression ^71^. However, the molecular mechanisms at GWAS loci associated with fatty acids in adipose tissue have not been fully understood. In particular, the epigenetics of fatty acids and the role of methylation of CpG sites in adipose fatty acid regulation are largely unexplored in adipose tissue. The effect of genetic polymorphisms on fatty acids is mediated through gene expression and DNA methylation ^71^ and methylation levels of CpG sites have been considered as regulators of gene expression levels ^73^. One experiment indicated that palmitate-treated human islets changed the global DNA methylation level linked with T2D ^74^. We integrated concurrent fatty acid and DNA methylation data in subcutaneous adipose tissue and observed the associations of arachidonic acid with three probes on chromosome 11 CpG island or shore, consistent with the previous study ^71^. We also identified that cg16744911 serves as the enhancer of *SCD* and is associated with palmitic acid level.

In the present study, eight novel variants were discovered to be associated with fatty acid levels or ratios in adipose tissue. SNPs at the 3p25.2 *locus* were associated with arachidonic acid level and the ratio of arachidonic acid/linoleic acid. This locus contains multiple plausible candidate causal genes, including *PPARG*, *MKRN2* and *TSEN2*. The transcription factor peroxisome proliferator-activated receptor (PPAR)-γ2 has been found to be expressed specifically for adipose tissue, where it plays a vital role in regulating adipogenic differentiation and lipid storage ^75^. *MKRN2* has been proven to regulate lipid profiling including total cholesterol levels ^76^ and we found that the secondary *MKRN2* eQTL was colocalized with the arachidonic acid GWAS signal. Moreover, we discovered the association between three CpG sites located at the *MKRN2* promoter and *TSEN2* enhancer or shelf and n-6 polyunsaturated fatty acids, suggesting the regulatory effects of CpG sites on adipose fatty acid. Interestingly, adipose arachidonic acid level and its conversion rate GWAS signals colocalized with kidney function GWAS signals, suggesting a link between genetic regulation of adipose arachidonic acid and renal health. An increase of metabolites including prostaglandins, thromboxane, and leukotrienes generated from arachidonic acid results in inflammatory damage to the kidney ^77^.

We also identified associations between the ratio of arachidonic acid to linoleic acid and genetic variants at the *ACSL3/MOGAT1 locus*. *ACSL3* is enriched in lipid droplets and belongs to a family of enzymes that convert free long-chain fatty acids into the substrates for lipid synthesis and β-oxidation, and fatty acyl-CoA esters ^78,79^; *MOGAT1* catalyzes the conversion of monoacylglycerols to diacylglycerols and is highly expressed in adipose tissue ^80^. An *in vitro* study showed PPARγ could regulate the transcription of *MOGAT1*, suggesting potential complex genetic regulatory pathways of n-6 polyunsaturated fatty acids ^81^. Other novel loci also play a vital role in the human body nevertheless do not have prior evidence of influencing fatty acid conversion in adipose tissue directly. *EXOC6B* encodes a protein required for exocytosis ^82^; *CYP26B1* encodes cytochrome-P450 enzymes that catabolize retinoic acid, which is essential for cell-cell signaling and cell growth ^83^ and the activity of *CYP26B1*may influence atherosclerosis ^84,85^; *ZBTB41/ASPM*, *LINC02055*, *MTBP*, *ADAMTSL1*, and *HAPLN1* have been implicated as biomarkers of cancer ^86–90^. However, we did not identify expressions of genes or DNA methylations at CpG sites mediating the genetic effects of novel loci on fatty acid conversions in adipose tissue at these loci.

Overall adiposity is highly correlated to molecular function and cell-type composition of adipose tissue. When investigating genetic regulation of obesity related traits, there is an ongoing debate on whether or not to adjust for BMI (or other measures of adiposity) or cell type composition in analyses. Adjusting for BMI will remove real biological effects that are mediated by BMI, and can also potentially introduce false positives via collider bias, as BMI is itself highly heritable ^91^. On the other hand, adjusting for BMI can improve power to detect associations that are not mediated by overall adiposity by removing noise. Given the advantage of both approaches, we chose to not adjust BMI in the main analyses as we wish to capture effects that may be mediated via BMI. However, we also performed sensitivity analyses by re-running the heritability estimates and GWAS analyses including the potential confounders, BMI and/or cell types, as covariates, and saw similar results across the sensitivity analyses (Table S3, Table S4 and Table S6).

Higher BMI is associated with an increase in body fat, which can alter the balance and composition of fatty acids in adipose tissue. We found that the polygenic score of BMI was negatively associated with saturated fatty acids but positively associated with unsaturated fatty acids. Obesity increases cardiovascular risk through risk factors such as increased fasting plasma triglycerides, high LDL cholesterol, low HDL cholesterol, elevated blood glucose and insulin levels and high blood pressure. ^92^Therefore, participants with a high BMI demonstrate a characteristic lipid profile marked by elevated triglycerides and reduced HDL. As expected, the triglycerides polygenic score showed a similar pattern of associations with fatty acid levels as BMI polygenic score, in contrast to polygenic scores of HDL and gluteofemoral adipose tissue. These results suggest that fatty acid levels are influenced by genetic risk of BMI, local adiposity, and lipid profile.

A limitation of this study is the relatively small sample size, so we are underpowered to detect more genome-wide significant loci of adipose fatty acids, particularly on chromosome X. Given the observed high heritability of adipose fatty acids, and number of suggestive hits in the GWASs, future studies in a larger population are warranted. Another limitation is that we only investigate female participants, therefore follow-up studies are needed to ensure the generality. Despite the multi-omics analyses, we can still integrate adipose fatty acids with other molecular layers, such as proteomics and microbiomics, which would provide complementary information to translate how the genome alters fatty acid metabolism in adipose tissue.

In summary, this study has provided a map of heritability and individual genetic variants that regulate fatty acid levels in adipose tissue. We identified genetic loci mediated through gene expression or DNA methylation in adipose tissue and linked the genetic regulation of fatty acids with biomedical phenotypes, thus helping to identify causal genes or CpG sites and providing pharmaceutical targets for metabolic diseases.

## Supporting information

TUK_Genetics_AdiposeFattyAcid_SupplementalMethodsFigures

TUK_Genetics_AdiposeFattyAcid_SupplementalTables

## Supplemental Data

Document S1. Supplemental Material and Methods, and Tables S1-S13.

Document S2. Figures S1-S10.

## Acknowledgements

This study was supported by an Accelerating Medicine Partnership Common Metabolic Disease award to K.S.S and J.T.B. K.S.S. acknowledges support from the MRC [MR/Y013891/1]. This work was in part supported by JPI HDHL-funded DIMENSION project (administered by the BBSRC UK, grant no. BB/S020845/1 to J.T.B.). X.Y. is funded by King’s-China Scholarship Council (K-CSC) joint scholarship. M.T. is funded by National Institute for Health and Care Research (NIHR) and Biomedical Research Centre (BRC). C.M. is supported by the Chronic Disease Research Foundation (CDRF) and by the Italian Ministry of Health - Bando Ricerca Corrente 2023. TwinsUK is funded by the Wellcome Trust, Medical Research Council, Versus Arthritis, European Union Horizon 2020, CDRF, Zoe Ltd and the NIHR Clinical Research Network (CRN) and Biomedical Research Centre based at Guy’s and St Thomas’ NHS Foundation Trust in partnership with King’s College London. The contribution of all participants of the TwinsUK is gratefully acknowledged. The authors acknowledge use of the research computing facility at King’s College London, King’s Computational Research, Engineering and Technology Environment (CREATE).

## Author contributions

X.Y. performed the main analyses and wrote the paper. K.S.S. designed and supervised the study. S.V. supervised by J.T.B. performed the follow-up meQTL analysis in TwinsUK. M.B.F. supervised GWAS analysis in TwinsUK. A.L.R. and J.S.E-S.M. contributed to genotype and gene expression data generation. M.T. retrieved and matched cardio-metabolic phenotypes. M.A-H. supervised by T.A.B.S. detected fatty acid proportions in adipose tissue in TwinsUK. C.M. contributed to generation and quality control of serum metabolite data. K.S.S. and J.T.B. contributed to supervision and overall curation and processing of the data. All authors reviewed and approved the manuscript.

## Declaration of Interests

The authors declare no competing interests.

## Data and code availability

Twins UK fatty acid GWAS datasets of 18 individual fatty acids (adjusted for age, BMI and family relatedness) can be accessed in the Knowledge Portal of Accelerating Medicines Partnership® Common Metabolic Diseases Consortium (AMP CMD Consortium) (https://hugeamp.org/dinspector.html?dataset=Small2023_FattyAcids_EU). Full GWAS summary statistics of all tested fatty acids are available within the CMD Knowledge Portal, allowing incorporation into PheWAS of other traits. TwinsUK fatty acid GWAS datasets for main analysis and sensitivity analysis are deposited in Zenodo (https://doi.org/10.5281/zenodo.15294670 and https://doi.org/10.5281/zenodo.15508762). TwinsUK RNA-Seq data are deposited in the European Genome-phenome Archive (EGA) under accession EGAS00001000805. TwinsUK methylation data are deposited in https://www.ebi.ac.uk/arrayexpress/experiments/E-MTAB-1866/. Access to TwinsUK individual level data can be obtained by application to the TwinsUK data access committee. For access information and how to apply, please refer to the website (https://twinsuk.ac.uk/researchers/access-data-and-samples/request-access/).

## Notes

https://hugeamp.org/dinspector.html?dataset=Small2023_FattyAcids_EU

https://doi.org/10.5281/zenodo.15294670

https://doi.org/10.5281/zenodo.15508762

https://ega-archive.org/datasets/EGAD00001001089

https://www.ebi.ac.uk/arrayexpress/experiments/E-MTAB-1866/

### Competing Interest Statement

The authors have declared no competing interest.

