## Supplementary material for "Genetic regulation of fatty acid content in adipose tissue": TUK_Genetics_AdiposeFattyAcid_SupplementalMethodsFigures

### Supplemental Material and Methods

#### Polygenic score

To construct polygenic scores (PGSs) for adiposity, BMI, waist-to-hip ratio, lipid biomarkers and chronic diseases, we searched terms on the PGS Catalog or GWAS performed in large cohorts of European ancestry. We calculated 13 PGSs: abdominal subcutaneous adipose tissue volumes adjusted for BMI (ASATadjBMI), visceral adipose tissue volumes adjusted for BMI (VATadjBMI), gluteofemoral adipose tissue volumes adjusted for BMI (GFATadjBMI), BMI, waist-to-hip ratio adjusted for BMI (WHRadjBMI), type 1 diabetes (T1D), T2D, coronary artery disease (CAD), hypertension, high-density lipoproteins cholesterol (HDL), low-density lipoproteins cholesterol (LDL), log-transformed total triglycerides (TG) and total cholesterol (TC) [1-7] (Table S1). This study analyzed the following 12 cardio-metabolic risk indicators: Percentage of fat in android region (AF), percentage of fat in the largest visceral fat region (VF), percentage of fat in gynoid region (GF), BMI, the ratio of percentage of fat in the largest visceral fat region to percentage of fat in gynoid region (VRGF), systolic blood pressure, diastolic blood pressure, HDL, LDL, TG, TC, and Atherosclerotic Cardiovascular Disease (ASCVD) risk score. AF, VAF and GF were measured via dual-energy X-ray absorptiometry scans (Hologic QDR; Hologic, Inc.) and VRGF was calculated afterward. During the clinical visit, the height and weight of the participants were measured by trained nurses following the harmonized protocols to calculate BMI. Blood pressure was measured using Marshall mb02, Omron Mx3 or Omron HEM713C Digital Blood Pressure Monitor with the participant sitting for at least three minutes by a trained nurse. Hypertension is considered to be from 140/90 mmHg or more (age < 80) or 150/90 mmHg or more (age ≥ 80) or self-reported anti-hypertension drug use. The lipid profiles and glycemic indicators levels were measured after an overnight fasting period according to standard procedures (Biochemistry Department, King's College Hospital, London and Affinity Biomarker Labs, London, United Kingdom). The 10-year ASCVD risk score is an algorithm utilized to estimate the cardiovascular risk of the participant. Detailed description of clinical indicators has been previously reported [8-10].

To validate the PGSs in TwinsUK, we hypothesized that the highest decile of PGS tends to reside in the highest decile trait or disease group, and vice versa. We only selected one twin in each pair and each trait was residualized against the covariates included in the original paper. We found that participants at the tails of the distribution for BMI and log triglycerides were enriched in extreme PGSs. For example, participants in the top 10% of the BMI distribution were nearly five times as likely to have a BMI polygenic score in the top 10% of the distribution (41.8% vs. 8.5%; OR = 4.92; 95% CI: 3.56 - 6.79). Conversely, individuals with less than the 10th percentile of BMI were over three times as likely to have a BMI polygenic score less than the 10th percentile (32.9% vs. 9.1%; OR = 3.61; 95% CI: 2.59 - 5.05) (Figure S1). The decile with the highest CVD PGS had a 1.73 (1.12 - 2.67) times higher risk of CAD.

To test the differences of morphological profiles on the tail ends of PGSs of diseases, individuals belonging to top 10% and bottom 10% of PGS distribution were categorized into two levels, then independent t-test was conducted. PGSs of CVD, T2D and hypertension revealed significant differences between disease and non-disease groups (t = -3.094, $P$ = 0.002; t = -4.235, $P = 2.4 \times 10^{-5}$; t = -5.378, $P = 8.3 \times 10^{-8}$). The adherence of PGSs to corresponding traits were predicted using the continuous PGS variable among all female twins adjusted for relatedness and covariates applied in the source paper. Overall, PGSs showed broadly strong associations with corresponding traits in TwinsUK cohort (Beta = 0.028 - 0.397, FDR < 0.05, Table S2).

### Supplemental Data

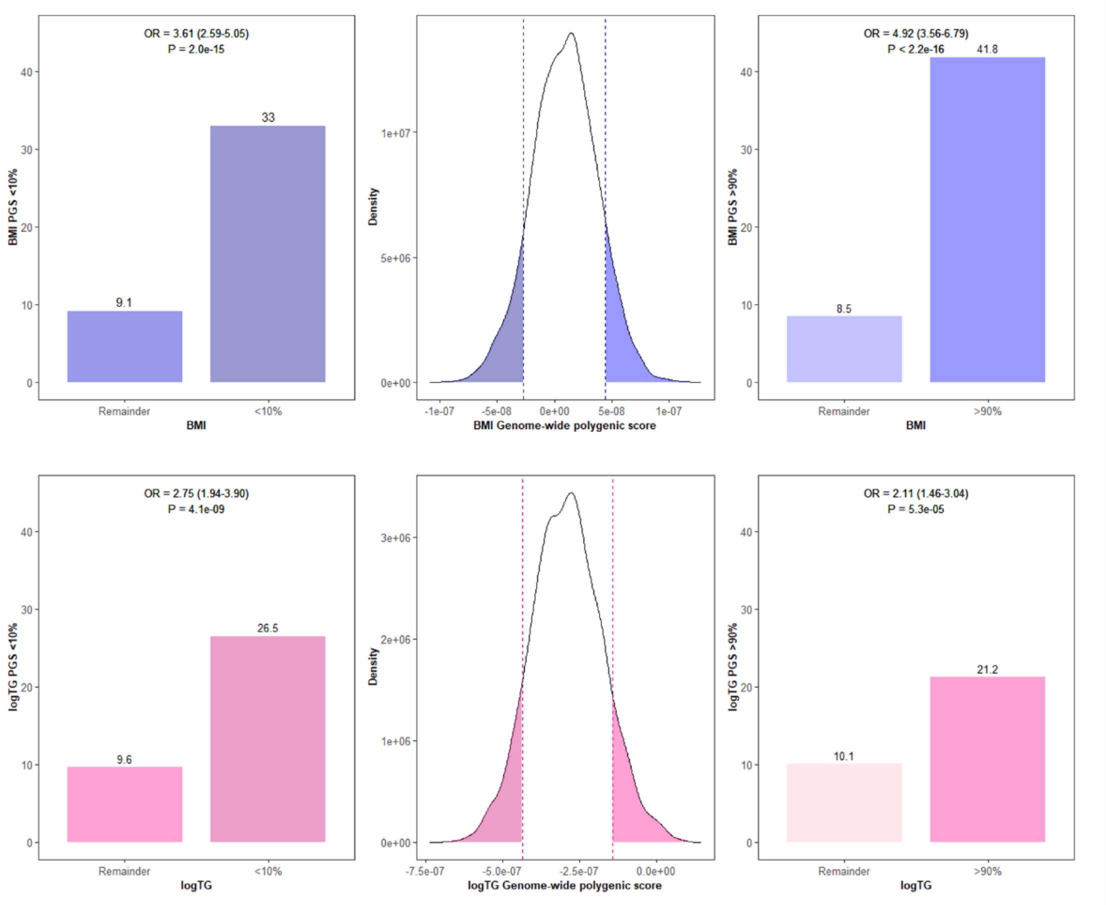


**Figure S1. Enrichment of BMI and log triglycerides genome-wide polygenic scores in tails of the distribution.**

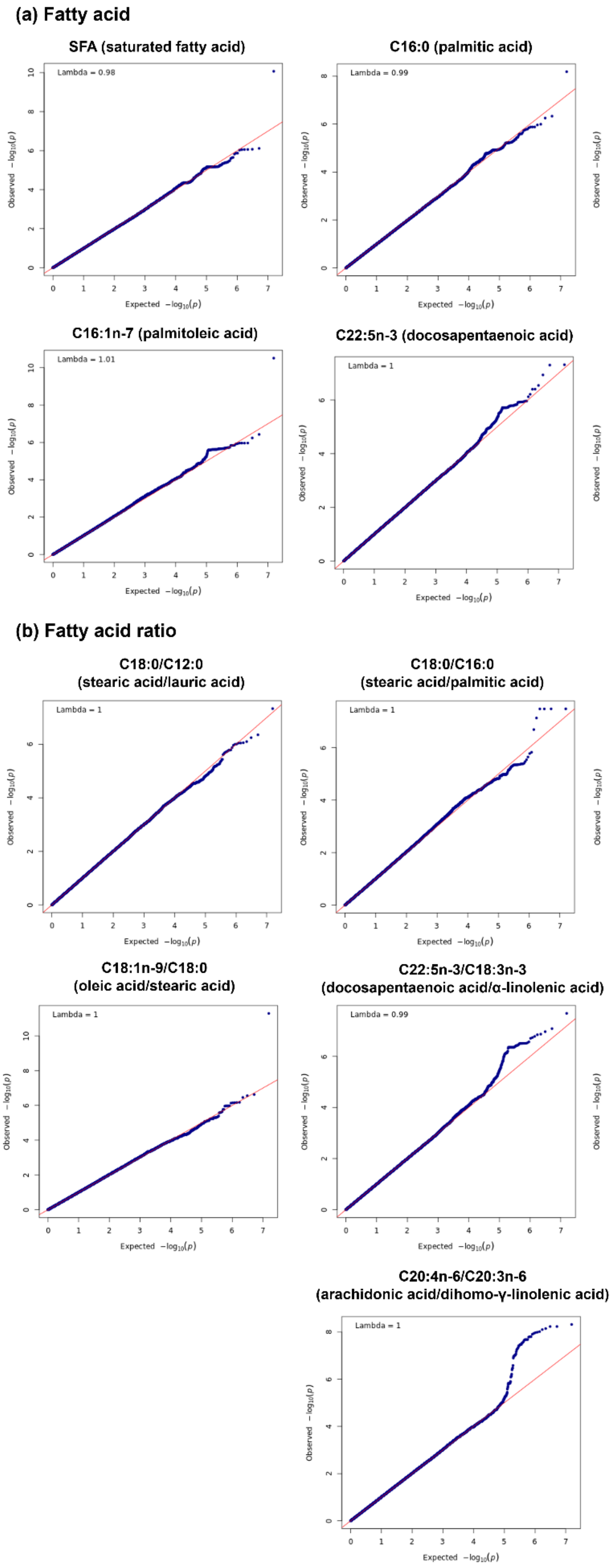


**Figure S2. QQ plots of GWASs on fatty acids in adipose tissue.**

QQ plots illustrate the findings of genome-wide association study for adipose fatty acids. The red diagonal line demarcates null hypothesis. Genomic inflation was close to one, suggesting no evidence of inflation.

#### (a) Fatty acid

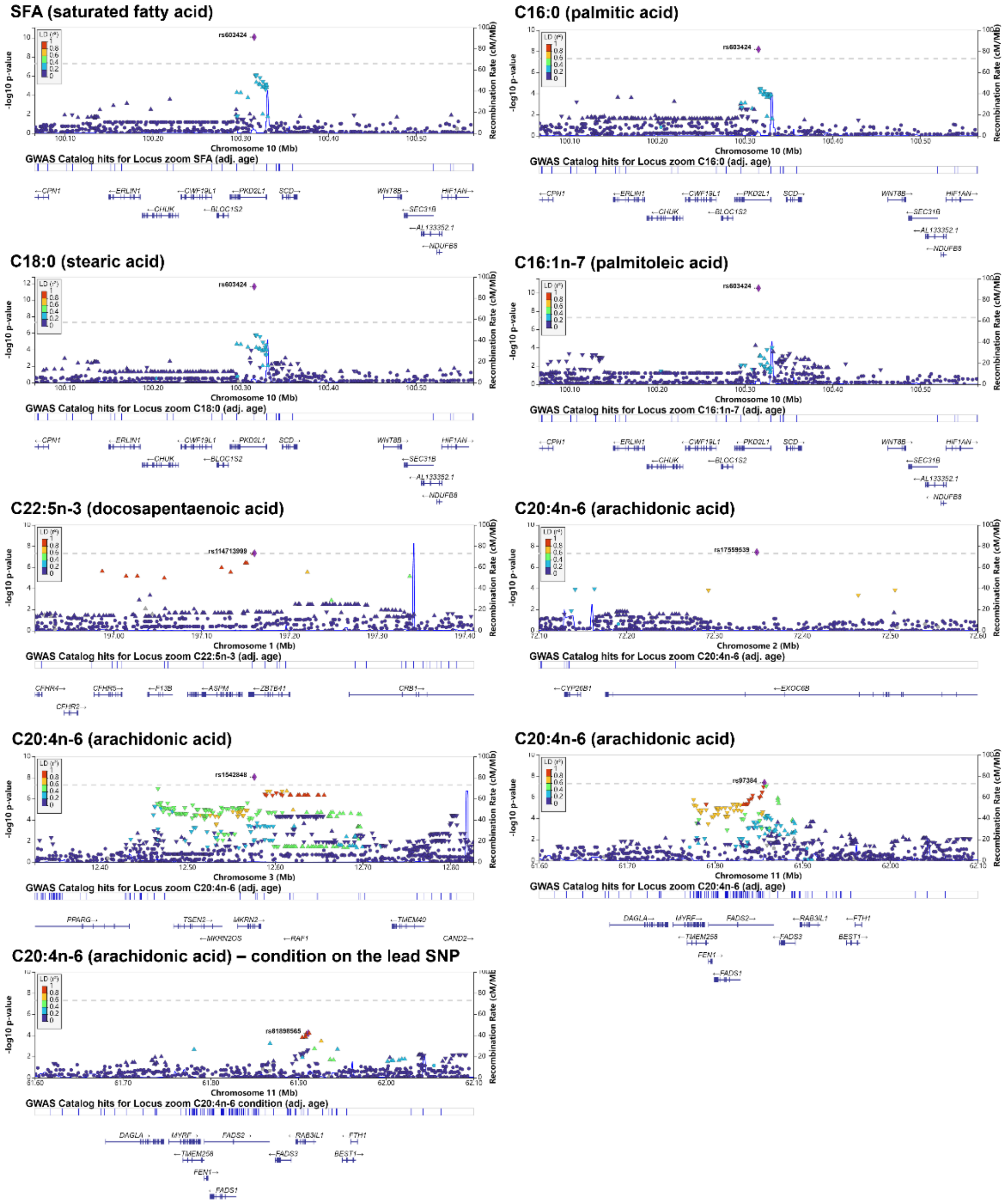
SFA (saturated fatty acid)
rs603424
Chromosome 10 (Mb)
GWAS Catalog hits for Locus zoom SFA (adj. age)
C16:0 (palmitic acid)
rs603424
Chromosome 10 (Mb)
GWAS Catalog hits for Locus zoom C16:0 (adj. age)
C18:0 (stearic acid)
rs603424
Chromosome 10 (Mb)
GWAS Catalog hits for Locus zoom C18:0 (adj. age)
C16:1n-7 (palmitoleic acid)
rs603424
Chromosome 10 (Mb)
GWAS Catalog hits for Locus zoom C16:1n-7 (adj. age)
C22:5n-3 (docosapentaenoic acid)
rs114713999
Chromosome 1 (Mb)
GWAS Catalog hits for Locus zoom C22:5n-3 (adj. age)
C20:4n-6 (arachidonic acid)
rs17559539
Chromosome 2 (Mb)
GWAS Catalog hits for Locus zoom C20:4n-6 (adj. age)
C20:4n-6 (arachidonic acid)
rs1542848
Chromosome 3 (Mb)
GWAS Catalog hits for Locus zoom C20:4n-6 (adj. age)
C20:4n-6 (arachidonic acid)
rs97384
Chromosome 11 (Mb)
GWAS Catalog hits for Locus zoom C20:4n-6 (adj. age)
C20:4n-6 (arachidonic acid) – condition on the lead SNP
rs61896565
Chromosome 11 (Mb)
GWAS Catalog hits for Locus zoom C20:4n-6 condition (adj. age)
-log10 p-value
Recombination Rate (cM/Mb)

#### (b) Fatty acid ratio

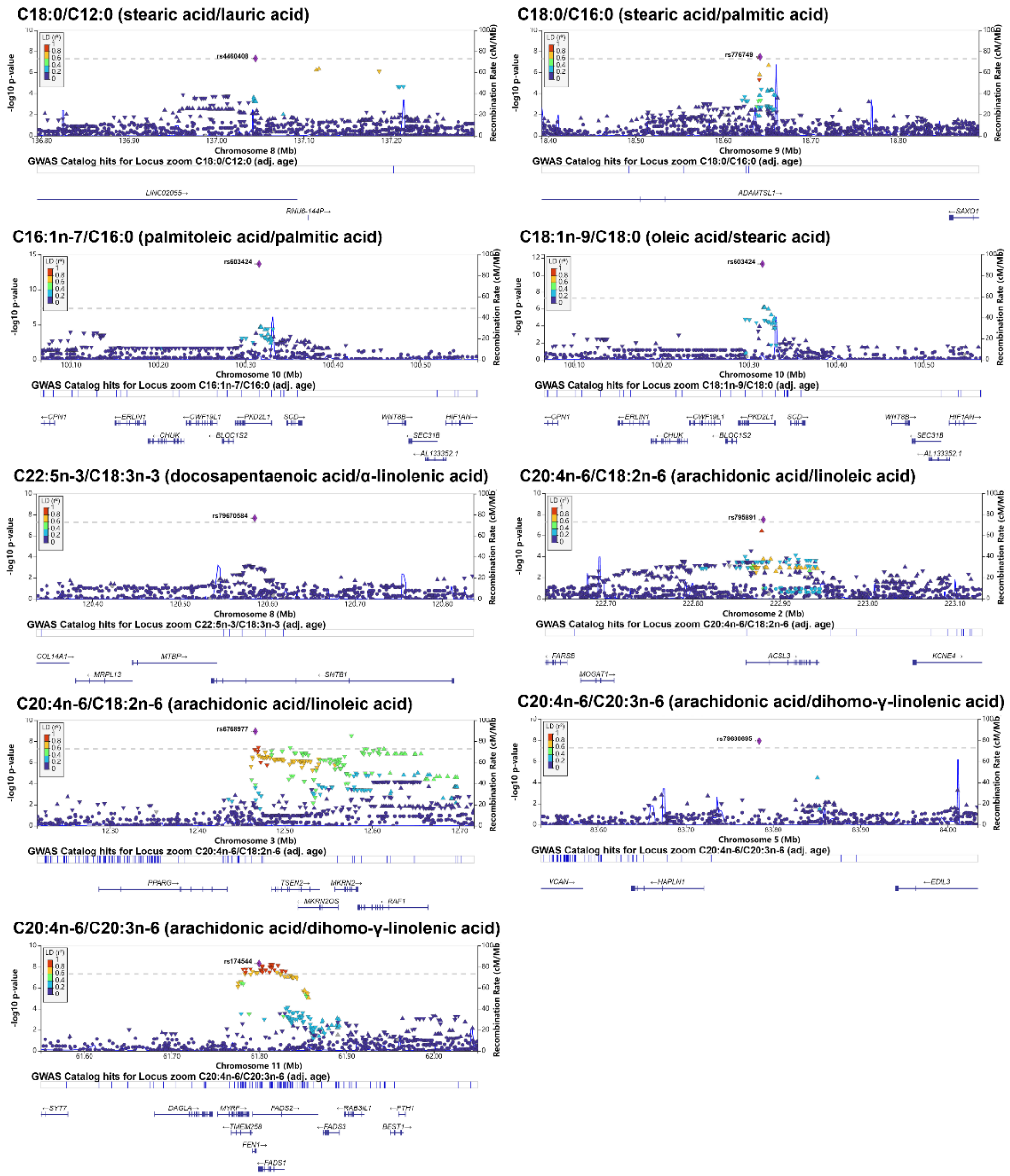


**Figure S3. Locus zoom plots of GWASs on fatty acids in adipose tissue.**

Locus zoom plots illustrate genome-wide association results of adipose fatty acids within several Mega base (Mb) pairs near significant loci.

#### (a) Fatty acid

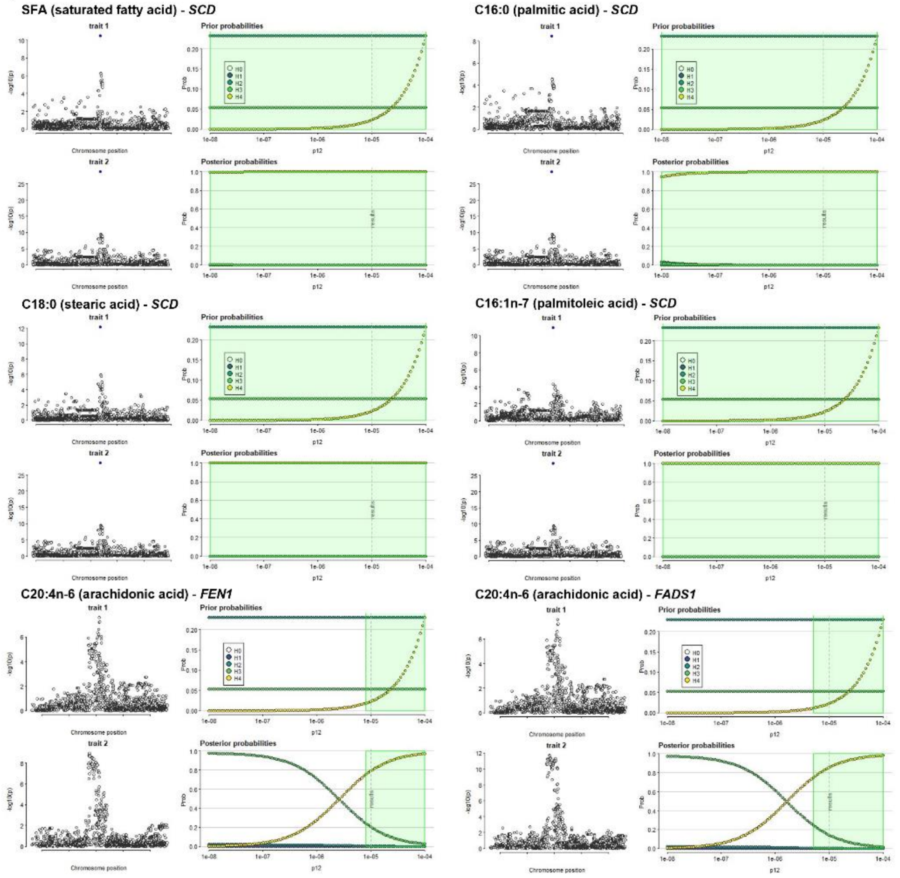


#### (b) Fatty acid ratio

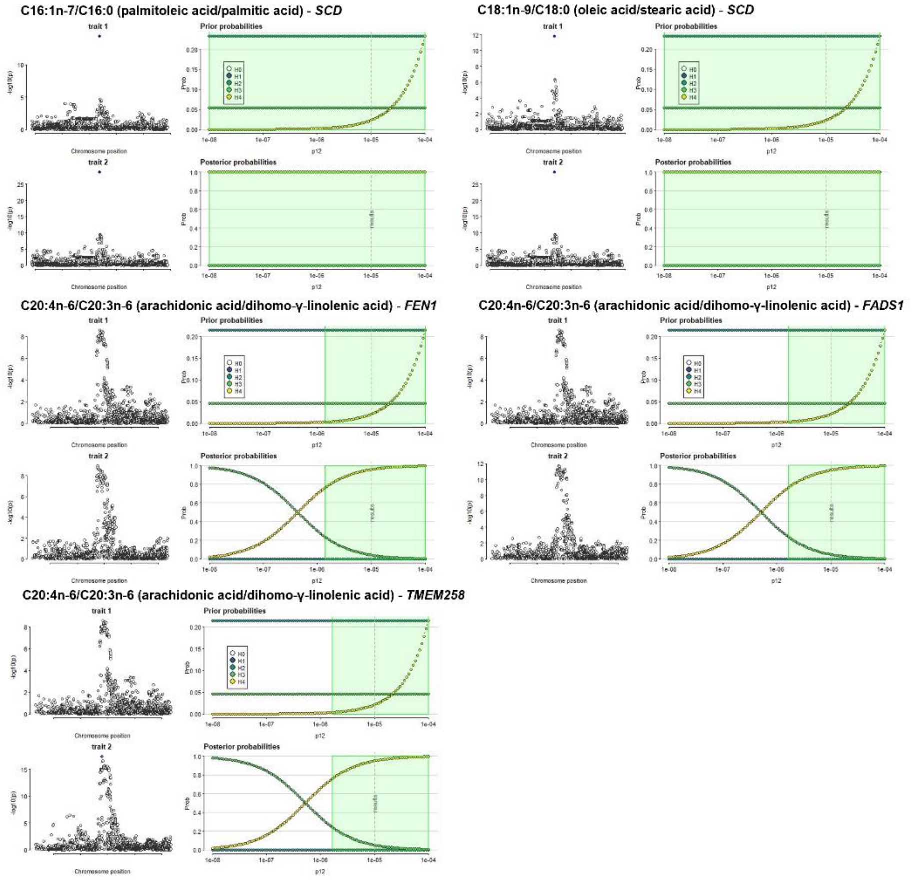

**Figure S4. Sensitivity analysis of colocalization between GWAS and eQTL signals.**

On the right the green region shows the region, the set of values of p12, for which PP4>0.75. On the left the input data are presented, with shading to indicate the posterior probabilities that a SNP is causal if H4 is true. In this case, the conclusion of colocalization looks quite robust.

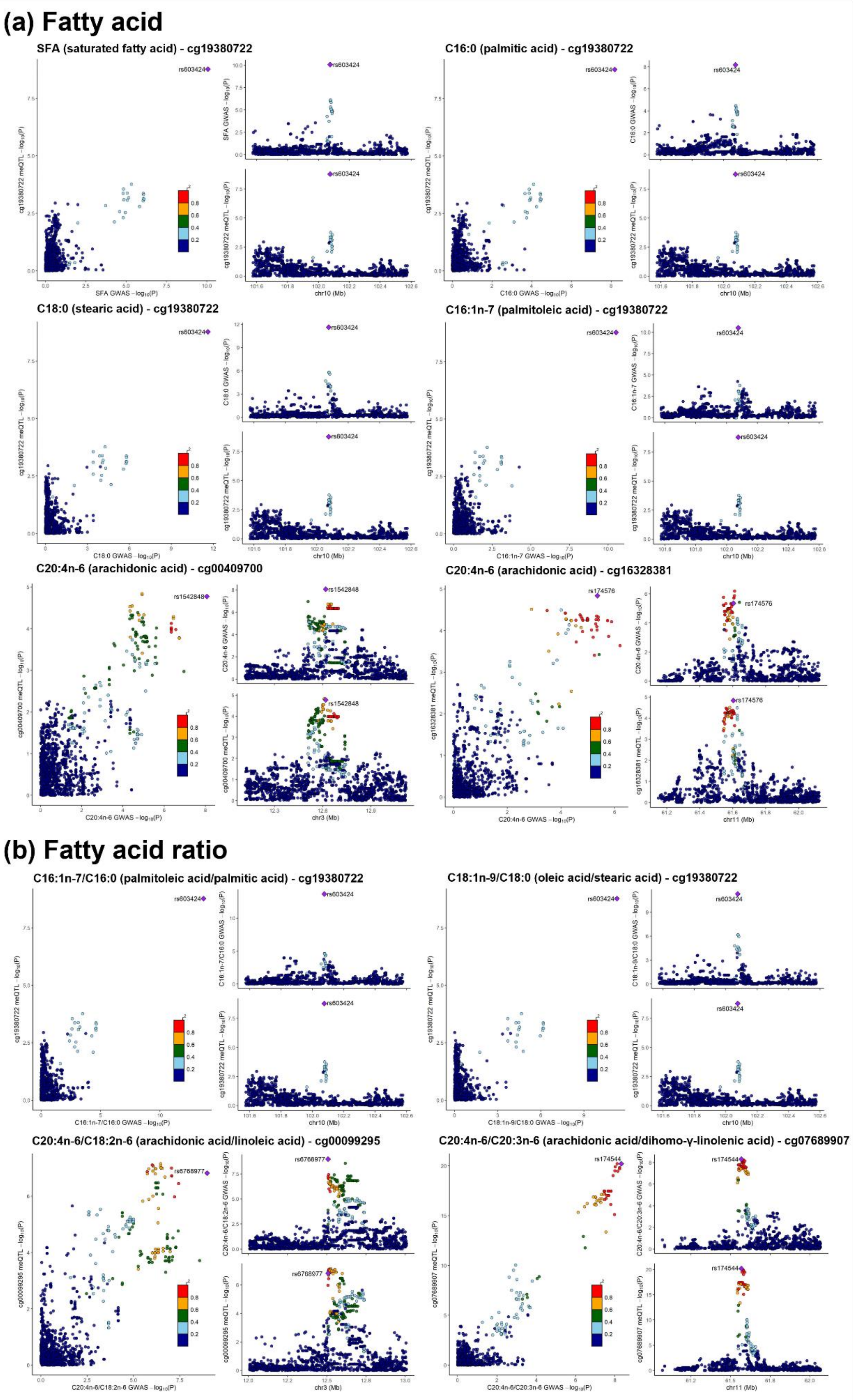


**Figure S5. Colocalization of GWAS and metQTL signals.**

The labeled SNP is the lead colocalized SNP for both GWAS and metQTL studies, and other SNPs are colored according to their LD $r^2$ with the lead SNP.

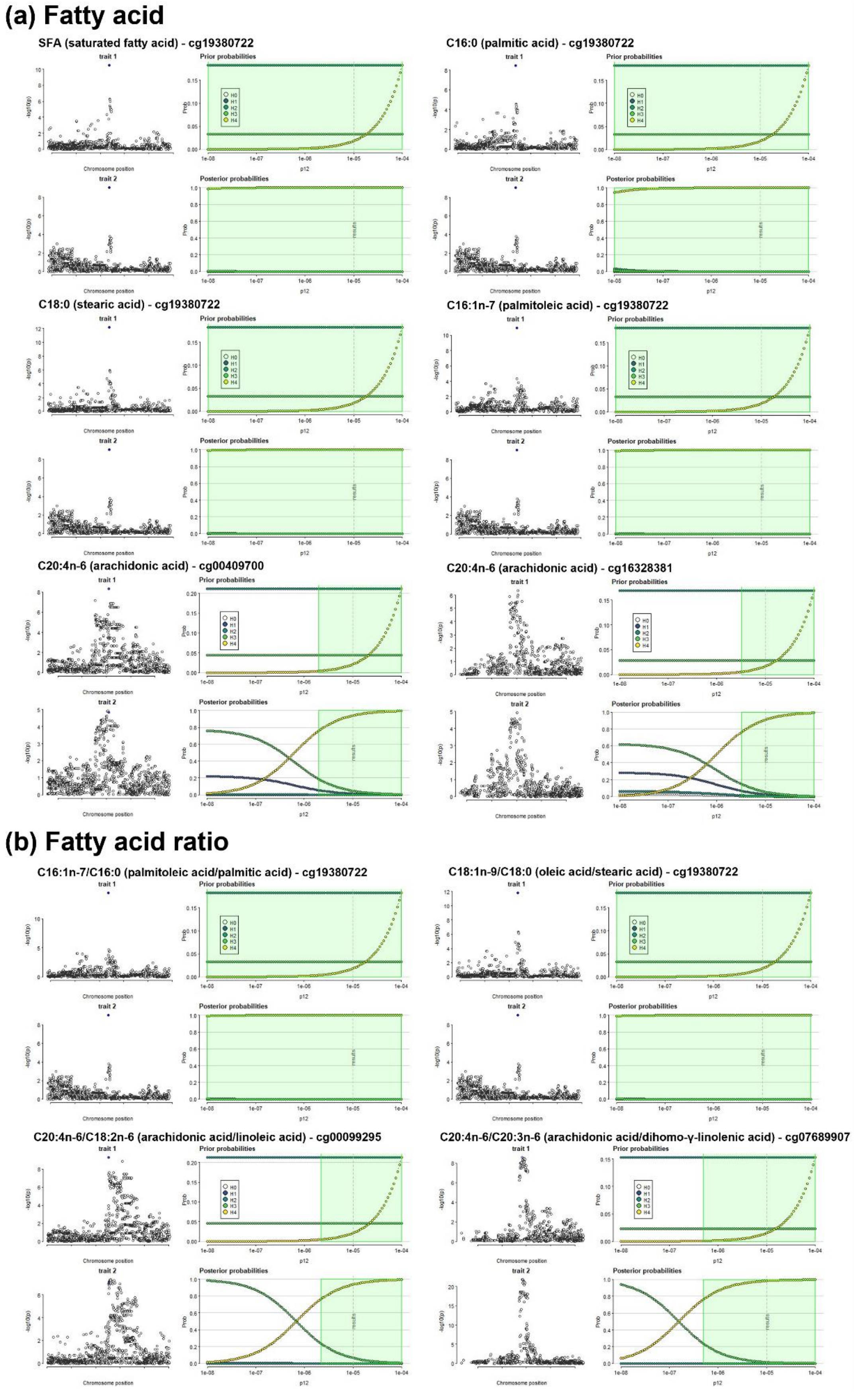


**Figure S6. Sensitivity analysis of colocalization between GWAS and metQTL signals.**

On the right the green region shows the region, the set of values of p12, for which PP4>0.75. On the left the input data are presented, with shading to indicate the posterior probabilities that a SNP is causal if H4 is true. In this case, the conclusion of colocalization looks quite robust.

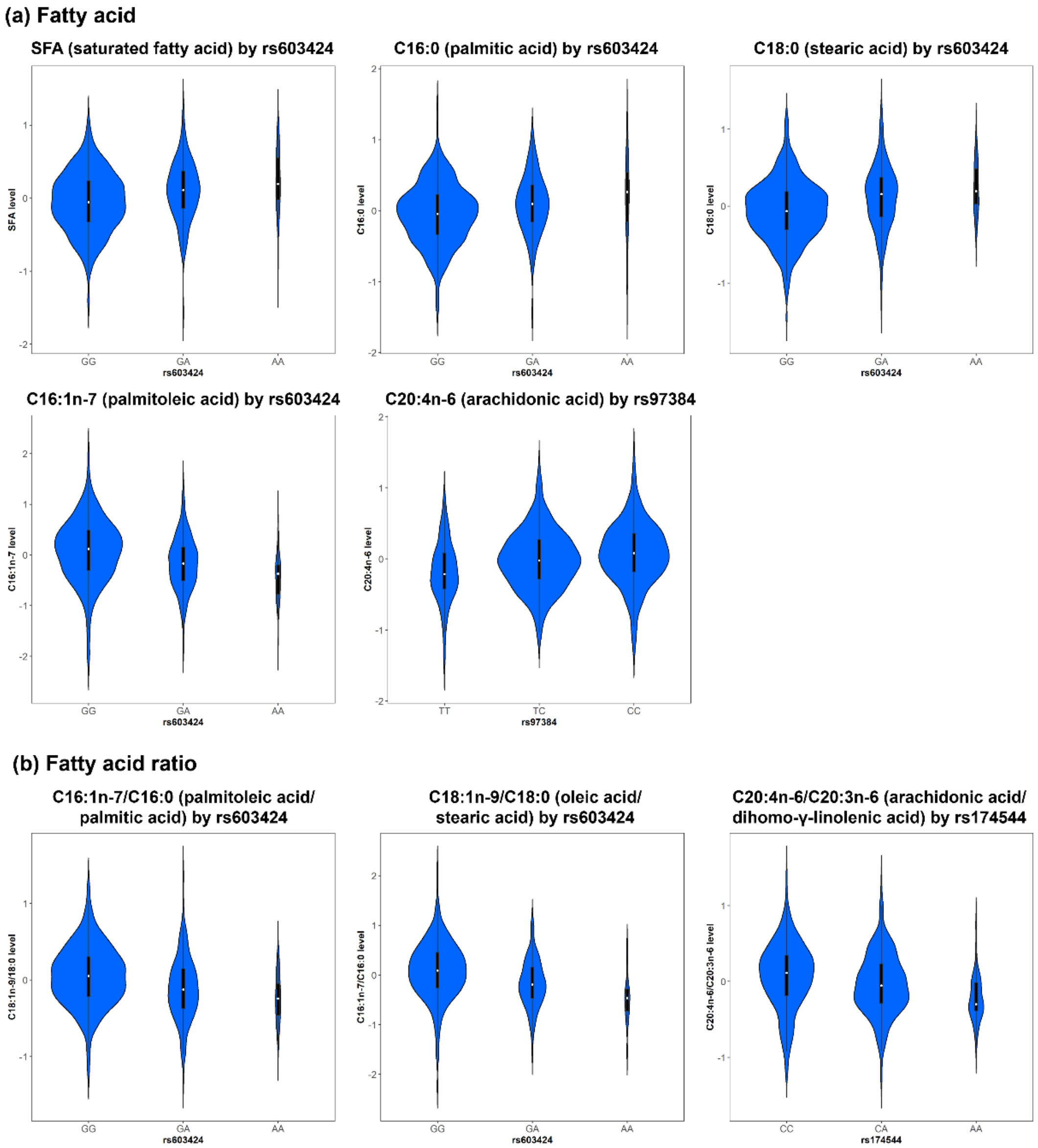


**Figure S7. Violin plots of fatty acids by genotypic class.**

The plots show the adiposee fatty acid levels stratified by genotypes i The effect of these SNP on fatty acids are potentially mediated through gene expression and/or DNA methylation.

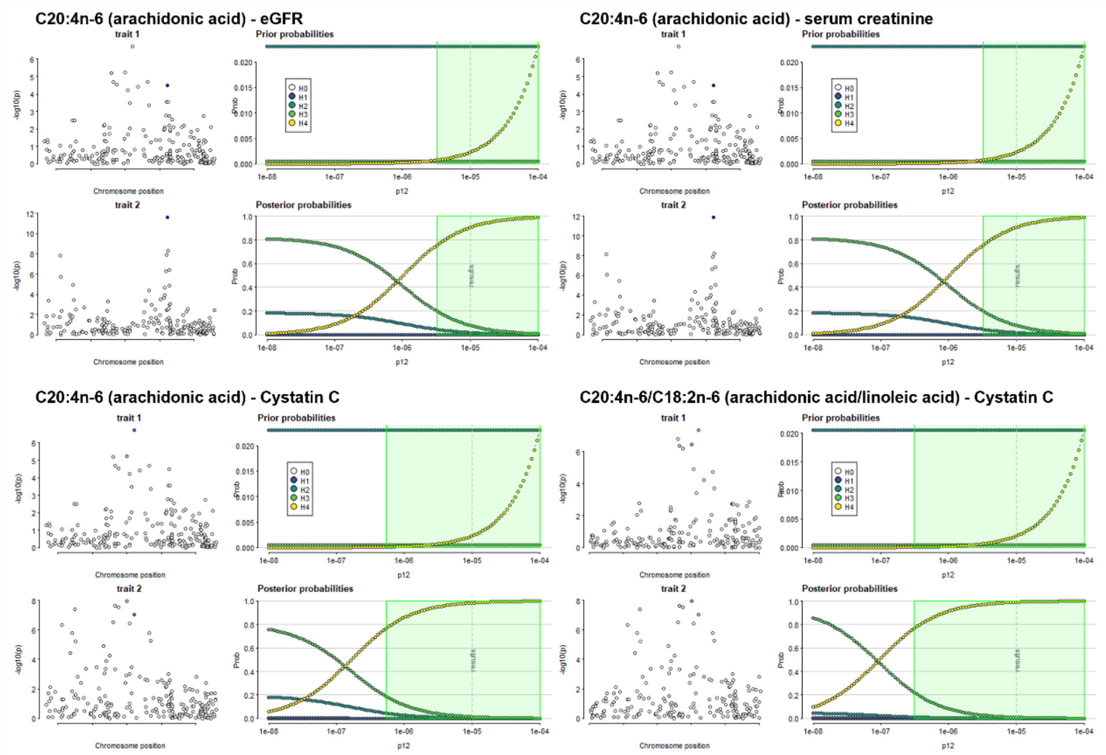


**Figure S8. Sensitivity analysis of colocalization between TwinsUK fatty acid GWAS and published kidney trait GWAS signals.**

On the right the green region shows the region, the set of values of p12, for which PP4>0.75. On the left the input data are presented, with shading to indicate the posterior probabilities that a SNP is causal if H4 is true. In this case, the conclusion of colocalization looks quite robust.

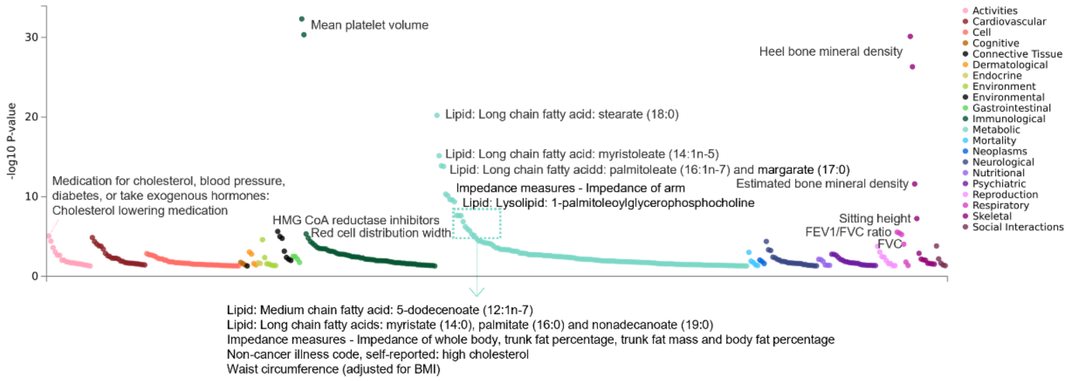


**Figure S9. Phenotypes associated with rs603424 at *SCD locus* (PheWAS plot based on GWASATLAS database).**

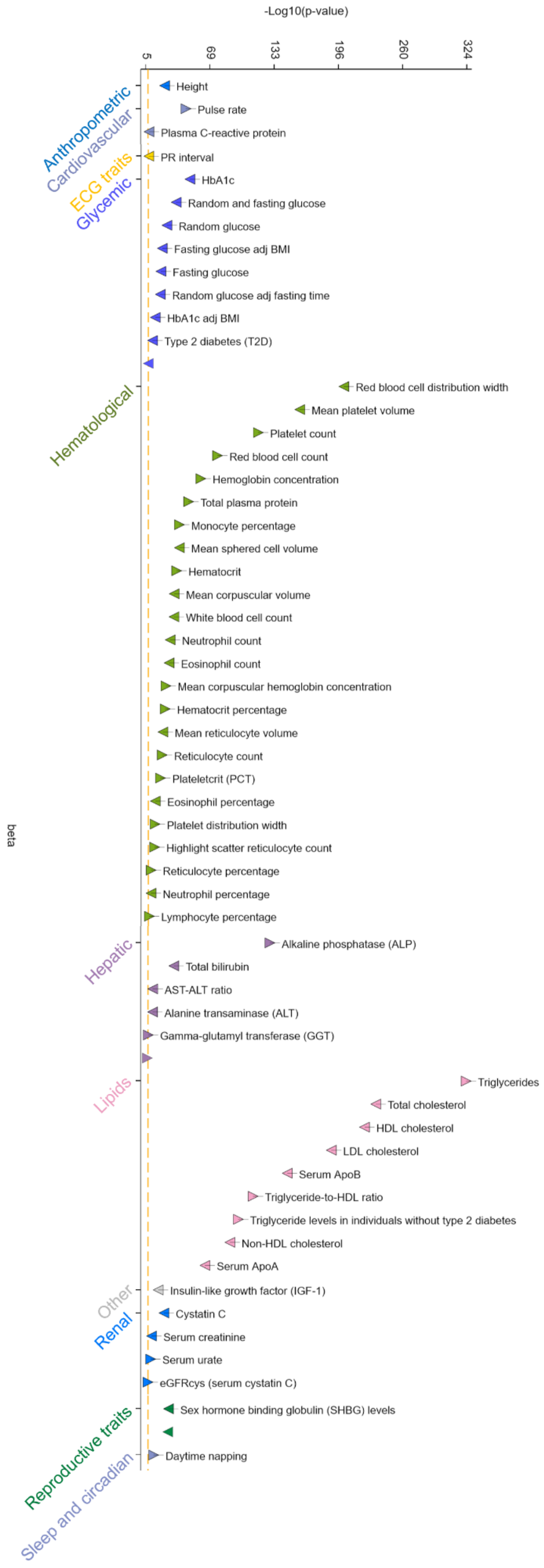


**Figure S10. Phenotypes associated with rs174544 at *FADS2 locus* (PheWAS plot based on AMP database).**
